# The *Trypanosoma brucei* subpellicular microtubule array is organized into functionally discrete subdomains defined by microtubule associated proteins

**DOI:** 10.1101/2020.11.09.375725

**Authors:** Amy N. Sinclair, Christine T. Huynh, Thomas E. Sladewski, Jenna L. Zuromski, Amanda E. Ruiz, Christopher L. de Graffenried

## Abstract

Microtubules are inherently dynamic cytoskeletal polymers whose length and activity can be altered to perform essential functions in eukaryotic cells, such as providing tracks for intracellular trafficking and forming the mitotic spindle. Microtubules can be bundled to create more stable structures that collectively propagate force, such as in the flagellar axoneme, which provides motility. The subpellicular microtubule array of the protist parasite *Trypanosoma brucei*, the causative agent of African sleeping sickness, is a remarkable example of a highly specialized microtubule bundle, comprising a single microtubule layer that is crosslinked to each other and the plasma membrane. The array microtubules appear to be highly stable and remain intact throughout the cell cycle, but very little is known about the pathways that tune microtubule properties in trypanosomatids. Here, we show that the subpellicular microtubule array is organized into subdomains that consist of differentially localized array-associated proteins. We characterize the localization and function of the array-associated protein PAVE1, which is a component of the inter-microtubule crosslinking fibrils present within the posterior subdomain. PAVE1 functions to stabilize these microtubules to produce the tapered cell posterior. PAVE1 and the newly identified PAVE2 form a complex that binds directly to the microtubule lattice. TbAIR9, which localizes to the entirety of the subpellicular array, is necessary for retaining PAVE1 within the posterior subdomain, and also maintains array-associated proteins in the middle and anterior subdomains of the array. The arrangement of proteins within the array is likely to tune the local properties of the array microtubules and create the asymmetric shape of the cell, which is essential for parasite viability.

**Author summary:** Many parasitic protists use arrays of microtubules that contact the inner leaflet of the plasma membrane, typically known as subpellicular microtubules, to shape their cells into forms that allow them to efficiently infect their hosts. While subpellicular arrays are found in a wide range of parasites, very little is known about how they are assembled and maintained. *Trypanosoma brucei*, which is the causative agent of human African trypanosomiasis, has an elaborate subpellicular array that produces the helical shape of the parasite, which is essential for its ability to move within crowded and viscous solutions. We have identified a series of proteins that have a range of localization patterns within the array, which suggests that the array is regulated by subdomains of array-associated proteins that may tune the local properties of the microtubules to suit the stresses found at different parts of the cell body. Among these proteins are the first known components of the inter-microtubule crosslinks that are thought to stabilize array microtubules, as well as a potential regulator of the array subdomains. These results establish a foundation to understand how subpellicular arrays are built, shaped, and maintained, which has not previously been appreciated.

## INTRODUCTION

*Trypanosoma brucei* is the causative agent of African trypanosomiasis, which affects both humans and livestock in Sub-Saharan Africa (1). A key contributor to *T. brucei* pathogenesis is the highly asymmetric shape of its cell body, which is essential for parasite survival within both insect and mammalian hosts. The trypomastigote form of *T. brucei* has a bore-like shape and tapered ends, with a broader cell posterior and a narrower, pointed anterior end (Figure 1A). This shape is produced by a single layer of microtubules that underlies the plasma membrane called the subpellicular array (2,3). A single flagellum is attached to the cell surface by the flagellum attachment zone (FAZ), which is comprised of a filament that is inserted between the subpellicular microtubules. The FAZ filament follows the left-handed helical path of the array microtubules as they traverse the cell body (4). As the flagellum beats, it deforms the subpellicular microtubule array and creates ‘cellular waveforms’ that define the distinctive corkscrew-like motility pattern of *T. brucei* (5-7). This specialized motility is essential for parasite escape from high-flow blood vessels and capillaries into low-flow compartments, which leads to the invasion of epithelial and adipose tissues that facilitate persistent host infection and dissemination back to the insect vector (8-11). The microtubules of the subpellicular array must be able to withstand and propagate the force generated by the flagellum, which is not uniformly distributed along the cell body (7). The ability to regulate microtubule dynamics and flexibility within different domains of the array would allow the parasite to optimize the transmission of energy generated by the flagellar beat and channel it into productive motility.

**Figure 1.**
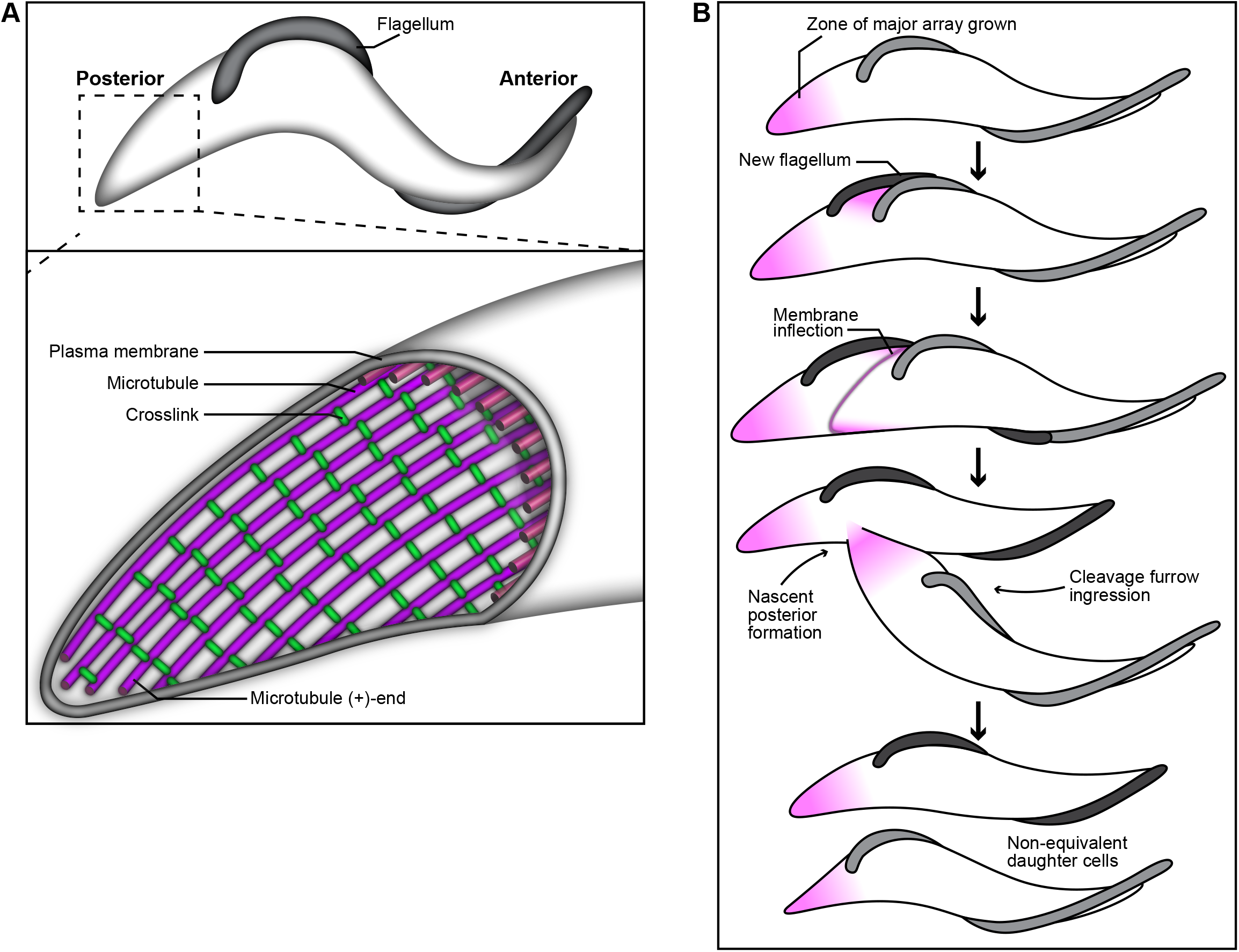
Schematics of the cytoskeleton and cell division in *Trypanosoma brucei*. **(A)** The subpellicular array is a single layer of microtubules purple) that underlies the plasma membrane. The microtubules are crosslinked to each other by regularly spaced fibrils (green). The (+)-growing ends of array microtubules face the cell posterior. **(B)** Array microtubules appear to be consistently polymerizing at the posterior end of the cell (purple shading). As the new flagellum is nucleated, new microtubules are inserted between the old and new flagellum to increase cell width. A membrane inflection forms that demarcates the path of cleavage furrow ingression, which follows the helical path of array microtubules. Cleavage furrow ingression initiates at the cell anterior, and nascent posterior end formation remodels the array at the cell midzone to resolve the subpellicular array into two distinct structures to complete cytokinesis.

Early morphological studies using electron microscopy revealed that the subpellicular array is a complex structure that is highly conserved among trypanosomatids (3,12). The microtubules of the array are uniformly oriented with the same polarity, with their growing plus-ends facing the broader cell posterior (13-15). The array microtubules are spaced apart at a uniform distance of 18-20 nm and are extensively crosslinked to each other and the plasma membrane by regularly spaced fibrils (16-18). The widest point of the cell body, which is near the nucleus, can contain over 100 microtubules. As the cell tapers toward the posterior and anterior ends, microtubules terminate in a stepwise fashion that maintains the inter-microtubule distance and consistent spacing between crosslinking fibrils while creating the asymmetric shape of the trypanosome cell body.

Microtubules are rigid, hollow polymers made up of α- and β-tubulin heterodimer subunits that bind and hydrolyze GTP. They exhibit a property known as dynamic instability wherein they undergo GTPase-dependent cycles of depolymerization (‘catastrophes’) and growth (‘rescues’) (19). This property allows for the rapid reorganization of internal cytoskeletal networks and is essential for cell migration, intracellular transport, and division (20). Microtubules are organized and regulated by microtubule associated proteins (MAPs) and motor proteins such as kinesins to control their dynamic instability. Many MAPs crosslink microtubules into bundles that can propagate and withstand force, such as the bundles found at the mitotic spindle, ciliary axoneme, and neuronal axon (21-24). Tubulin subunits have intrinsically disordered and highly charged C-terminal tails that are heavily decorated with post-translational modifications, which serve as binding sites for many MAPs (25,26). Other MAPs recognize specific structural conformations of the microtubule lattice, such as the incompletely polymerized and more open lattice found at microtubule plus-ends (27,28).

The subpellicular microtubule array is a fascinating example of a specialized microtubule bundle. Dynamic instability has not been observed in microtubules of the array, most likely due to their extensive crosslinking (Fig. 1A). Some crosslinking proteins in other systems are known to promote microtubule flexibility (29), which may explain the apparent longevity of array microtubules as they respond to forces produced by the flagellar beat. However, very little is known about the structure and function of the inter-microtubule crosslinking fibrils that organize the array, or how MAPs may regulate array microtubules. Fewer than twenty non-motor proteins have been identified that associate with the subpellicular array (30). The function of many of these has yet to be determined, but a subset have been described as microtubule stabilizers or potential crosslinkers that are required for proper cell division (31-34) and the arrangement of array microtubules in a single layer underneath the plasma membrane (35-37). However, the precise mechanisms these proteins employ to localize to the array and perform their function have not been established. Understanding how *T. brucei* MAPs bind to microtubules will have important implications in trypanosomatid cell biology and provide a more fundamental understanding of how cytoskeletal filaments are organized among diverse phyla.

The array microtubules appear highly stable and remain intact throughout the cell cycle, which requires that many aspects of *T. brucei* cell division include microtubule remodeling processes that duplicate and segregate the array (13,38). During cell division, extant microtubules are elongated and short, new microtubules are inserted between them to lengthen and widen the array (16). At later stages of the cell cycle, new microtubules are inserted in between the old and new flagellum to widen the cell body (Fig. 1B) (39). Microtubule polymerization also appears to occur continuously at the cell posterior throughout the cell cycle, suggesting that the array microtubules may be consistently turning over tubulin subunits to maintain cell length and shape (14,39,40). Cytokinesis proceeds via a cleavage furrow that initiates at the cell anterior and follows the helical path of the array microtubules towards the cell posterior to divide the cell on its long axis (Fig. 2B) (13,38,41).

**Figure 2.**
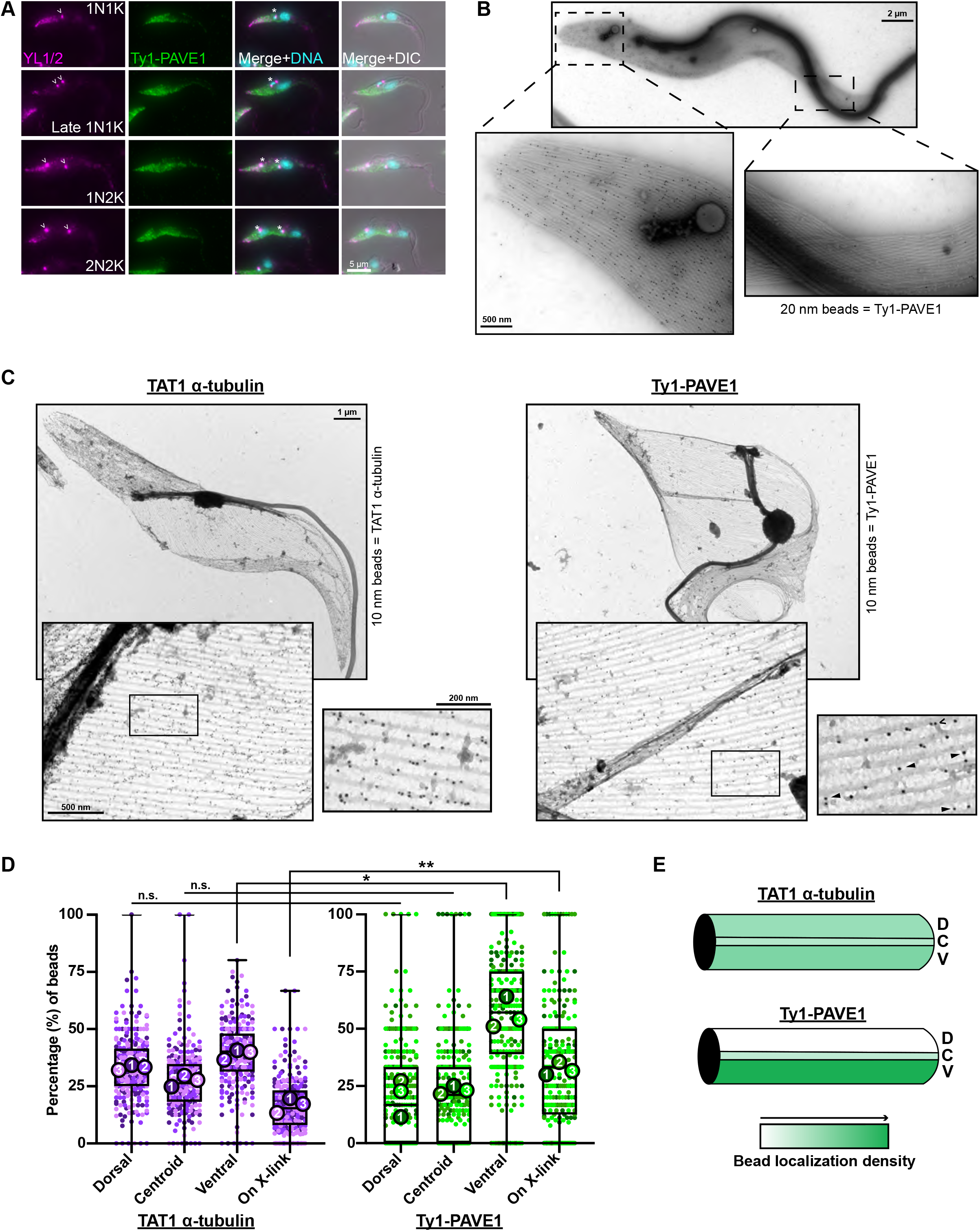
PAVE1 is a component of the inter-microtubule crosslinks of the cell posterior. **(A)** Ty1-PAVE1 cells were fixed in methanol and labeled with anti-Ty1 antibody and YL1/2, which recognizes tyrosinated tubulin. The empty arrowheads are denoting pools of tyrosinated tubulin at the basal body. Asterisks are marking kinetoplasts. **(B)** Whole-mount extracted cytoskeleton immunogold-labeled for Ty1-PAVE1 with 20 nm beads. **(C)** (Left) A subpellicular array sheet from WT 427 control cells immunogold-labeled with 10 nm beads against TAT1, which recognizes trypanosome a-tubulin. (Right) A subpellicular array sheet immunogold-labeled for Ty1-PAVE1 with 10 nm beads. Empty arrowheads are pointing to ventrally localized Ty1-PAVE1 beads, and solid arrows are marking Ty1-PAVE1 beads on preserved inter-microtubule crosslinking fibrils. **(D)** Quantification of immunogold label distribution along individual microtubules in subpellicular sheets represented as a Superplot (73), which shows each individual measurement in each biological replicate. The replicates are color-coded and the average is illustrated as large circles. TAT1 a-tubulin—Ty1-PAVE1 dorsal:dorsal and centroid:centroid distributions are not significantly different (P=0.0542 and P=0.0741, respectively). *P=0.0146, **P=0.0032. P-values are calculated from unpaired two-tailed Student’s *t-*tests using the averages from N=3 independent biological replicates. 3 subpellicular sheets in both TAT1 control and Ty1-PAVE1 conditions from N=3 independent biological replicates were quantified. (E) Heat map of a-tubulin and Ty1-PAVE1 immunogold label distribution on individual microtubules.

In this work, we have identified a set of *T. brucei* proteins that localize to different subdomains of the subpellicular array. These proteins play important roles in maintaining the shape of the array and the composition of these subdomains. We define the function of an array-associated protein known as PAVE1, which is found exclusively in kinetoplastids and localizes to the posterior and ventral edge of the array (42). We previously showed that PAVE1 is required for the formation of the more broadly tapered end of the cell posterior. We now show that PAVE1 is a component of the inter-microtubule crosslinking fibrils present in the posterior subdomain of the array and is essential for stabilizing the growing microtubules found within this subdomain. PAVE1, in complex with its binding partner PAVE2, binds directly to the microtubule lattice, demonstrating that PAVE1 and PAVE2 together form a true kinetoplastid-specific MAP. The previously identified protein TbAIR9 (34) forms a complex with PAVE1 and PAVE2 and regulates the distribution of array-associated proteins to specific subdomains of the subpellicular array. Our work shows that the subpellicular array contains a landscape of differentially localized array-associated proteins that likely function to locally tune the biophysical characteristics of array microtubules. These results update the long-standing view of the subpellicular array as a static arrangement of microtubules. Our work suggests that the array is a dynamic, multidomain structure that plays an essential role in cell morphogenesis and motility.

## RESULTS

### PAVE1 localizes to the inter-microtubule crosslinks of the subpellicular array at the cell posterior

PAVE1 (Posterior And Ventral Edge protein 1) is an array-associated protein that localizes to the posterior and ventral edge of the subpellicular array (42). To determine if PAVE1 co-localizes to regions of new microtubule growth, we compared PAVE1 distribution to the labelling pattern of the antibody YL1/2, which recognizes the terminal tyrosine residue of alpha-tubulin (43). Terminal tyrosination is a hallmark of newly polymerized tubulin and is considered a marker of array growth during the cell cycle (16,39,44). We generated a cell line containing an endogenously tagged PAVE1 with three copies of the Ty1-epitope tag at its N-terminus, stained the cells with anti-Ty1 and YL1/2 antibodies, and examined their distribution throughout the cell cycle. Trypanosomes contain one nucleus (N) and one kinetoplast (K; the mitochondrial DNA aggregate) early in the cell cycle (1N1K). Cells first duplicate their kinetoplast (1N2K) before undergoing nuclear mitosis (2N2K) prior to initiating cytokinesis. Ty1-PAVE1 and YL1/2 both stain the posterior array of cells early in the cell cycle (1N1K) with the labeling density declining as the array widens near the location of the kinetoplast (Figure 2A, asterisk). YL1/2 also labels a pool of tyrosinated tubulin present at the basal body (Fig. 2A, arrows). In late 1N1K cells, which can be identified by their elongated kinetoplasts, the YL1/2 signal declines at the cell posterior while PAVE1 signal remains constant. In 1N2K cells, YL1/2 signal increases and becomes more concentrated at the cell posterior, indicating more localized array growth, with no change in PAVE1 labeling pattern. In 2N2K cells, the PAVE1 signal extends past the midzone of the cell towards the more anterior-located nucleus, which is the location of extensive microtubule remodeling that leads to the creation of the new posterior end during cytokinesis (39). This localization pattern suggests that PAVE1 is stably associated with the array microtubules at the posterior end of the cell throughout the cell cycle and is recruited to the nascent posterior end during its formation.

PAVE1 may localize to specific regions of individual microtubules in the subpellicular array to carry out its function. We performed negative-stain immunogold electron microscopy (iEM) on extracted cytoskeletons prepared from cells containing Ty1-tagged PAVE1 to obtain high-resolution localization information (Fig. 2B). The 20 nm gold particles labeling Ty1-PAVE1 localized to the posterior portion of the subpellicular array with very little labeling on the cell anterior. However, it was not possible to determine if PAVE1 preferentially bound a specific region of individual microtubules. When whole-mount cytoskeletons are imaged by TEM, both the top and bottom layers of the array are visible, which results in a cross-hatching pattern that makes it difficult to distinguish individual microtubules. To address this problem, we generated subpellicular array sheets from immunogold-labeled cytoskeletons (16). Extracted whole-mount immunogold-labeled cytoskeletons were positively stained with uranyl acetate, critical point dried, and cleaved using double-sided tape to ‘unroof’ the cell and remove the top layer of the subpellicular array (Fig. 2C).

To determine if PAVE1 had a unique and polarized labeling pattern along individual microtubules, we compared PAVE1 distribution to the localization pattern of the TAT1 antibody, which recognizes *T. brucei* a-tubulin (Fig. 2C, left). We developed a semi-automated macro in ImageJ that allowed us to quantify the localization of immunogold labeling (Supplementary Fig. 1). Subpellicular sheets were oriented along their dorsal-ventral axis using the intact flagellum as a fiducial marker. Gold beads were then classified as localizing to the dorsal, centroid, or ventral regions of individual array microtubules. We found that in comparison to TAT1 labeling, PAVE1 preferentially localized to the outside ventral walls of the array microtubules at the cell posterior, as well as to the inter-microtubule crosslinks that remained after sheet preparation (Fig. 2D & E). Many crosslinks were disrupted in the sheet preparation, leaving small protrusions on the outside microtubule wall (Fig. 2C, open arrowhead). The ventral localizations of PAVE1 often occurred on these protrusions, likely indicating that PAVE1 was on a crosslink that was disrupted. This localization pattern suggests that PAVE1 may be part of the inter-microtubule crosslinking fibrils present in the posterior subdomain of the array.

### PAVE1 maintains microtubule length at the posterior subpellicular array

Nothing is currently known about how the microtubule crosslinking fibrils are assembled or remodeled as new tubulin dimers are added to the plus-ends present at the cell posterior. To determine how PAVE1 and the inter-microtubule crosslinks are incorporated into the array, we developed a capping assay using HaloTag technology, which allows the irreversible conjugation of a dye to a specific fusion protein (45). We appended the HaloTag protein to the N-terminus of one of the endogenous PAVE1 alleles, then incubated an exponentially growing culture with a cell-permeable HaloTag substrate containing the far-red fluor JF646 to covalently label all the HaloTag-PAVE1 present in the cell. The remaining JF646-conjugated substrate was then washed out of the culture and replaced with a substrate linked to the rhodamine analog TMR, which is only able to label newly synthesized HaloTag-PAVE1 (Fig. 3A). Cells were then harvested, fixed, and imaged using epifluorescence microscopy. The fluorescence intensity of each fluor-conjugated version of PAVE1 was analyzed using a line scan placed along the ventral edge of the cell body of 1N1K cells (Fig. 3B). This two-color labeling approach allowed us to localize where newly synthesized PAVE1 was added to the array and how its distribution changed over time. We designated T=0 h as the point when the TMR substrate was added to the media for our subsequent analysis.

**Figure 3.**
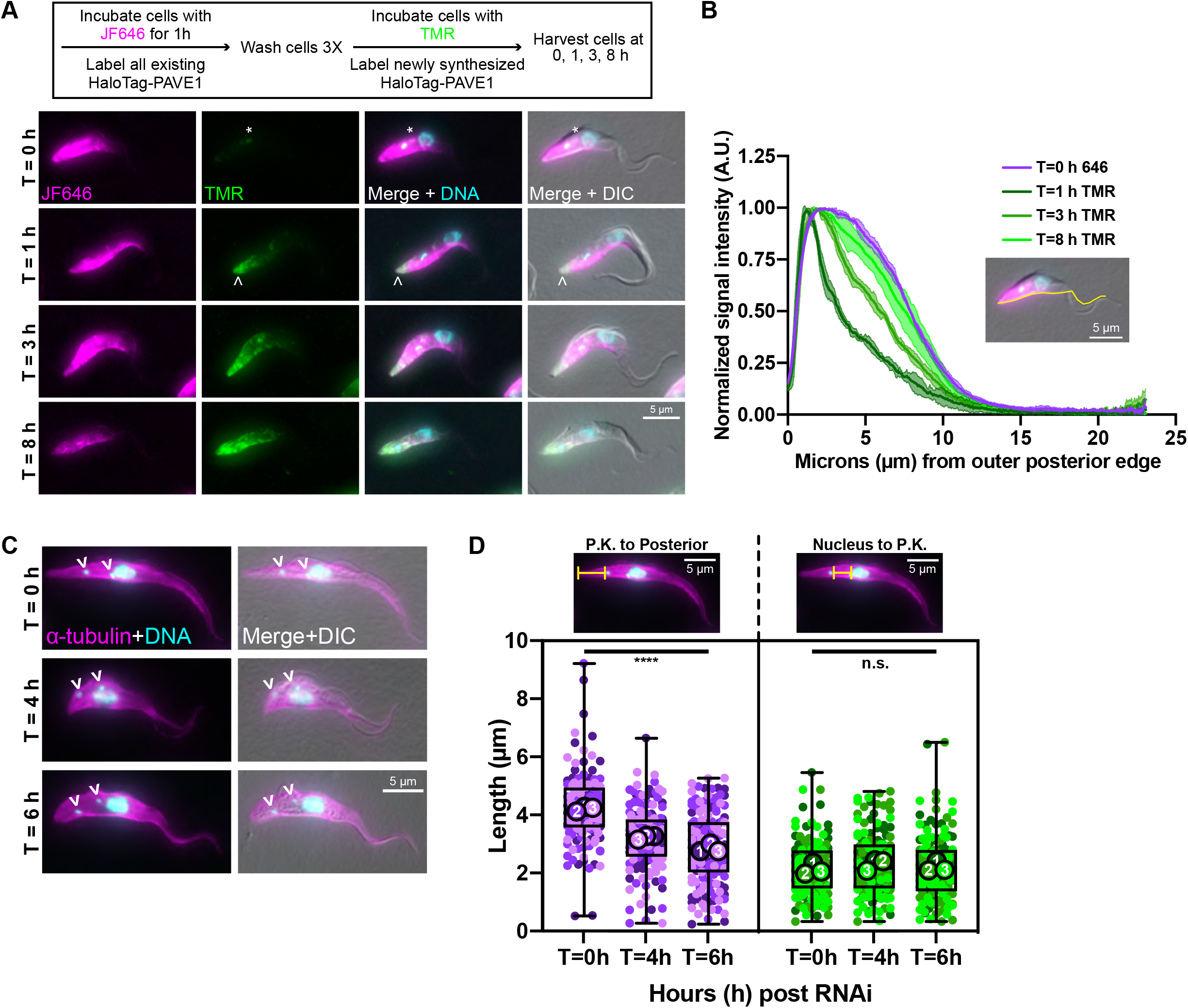
PAVE1 is added to and stabilizes the microtubules at the extreme posterior end of the array. **(A)** HaloTag-PAVE1 cells were fixed in paraformaldehyde at T=0 h, T=1 h, T=3 h, and T=8 h after the addition of HaloTag ligand conjugated to TMR to cell culture in the capping assay. Asterisk represents background labeling of the endomembrane system. Empty arrowhead is pointing to the earliest addition of HaloTag-PAVE1, represented by TMR labeling, at the extreme cell posterior. **(B)** Fluorescence intensity quantification of HaloTag ligand labeling along the ventral edge of the cell body at T=0 h, T=1 h, T=3 h, and T=8 h after TMR substrate addition. Fluorescence intensity was normalized to 1 at each time point in order to illustrate the shape of the curve. 50 1N1K cells at each time point from N=3 independent biological replicates were measured. **(C)** PAVE1 RNAi was induced and cells were harvested, fixed in methanol, and stained with anti-a-tubulin after T=0 h, T=4 h, and T=6 h of PAVE1 RNAi induction. 1N2K cells were selected for analysis as they completed cytokinesis prior to PAVE1 depletion. **(D)** Superplot of the measured distances between the posterior kinetoplast to outer posterior edge of the subpellicular array, and the posterior edge of the nucleus to the posterior kinetoplast in control and PAVE1 RNAi cells. One-way ANOVAs were calculated using the averages from N=3 independent biological replicates. ****P<0.0001, n.s. P=0.5653. 50 1N2K cells were measured at each time point from N=3 independent biological replicates.

At T=0 h, the JF646 signal labeled the posterior and ventral edge of cells in a similar pattern to what was observed with Ty1-PAVE1 immunofluorescence. There was an additional punctum of JF646 and TMR signal near the endomembrane system (Fig. 3A, asterisk), which was likely due to endocytosis of the HaloTag substrate. At T=1 h, the TMR signal was greatest at the extreme posterior end of the cell and rapidly declined towards the cell anterior (Fig. 3B). This suggested that new HaloTag-PAVE1 was added to the posterior subpellicular array at the same location that microtubule polymerization occurs throughout the cell cycle (14,16). At T=3 h, the TMR signal began to increase past the extreme posterior end. This demonstrated that HaloTag-PAVE1 moved toward the cell anterior over time. At the latest time point T=8 h, which was approximately the duration of one cell cycle, the TMR signal was similar to the original JF646 signal. Moreover, the JF646 signal intensity declined at T=8 while the TMR signal intensity increased, which suggested that there was turnover of HaloTag-PAVE1 on the array.

Depletion of PAVE1 using RNA interference (RNAi) truncates the subpellicular array until the posterior cell edge directly abuts the kinetoplast (42). We performed an RNAi time course to test if PAVE1 is required to maintain the microtubules of the posterior array, or if the array truncation seen in PAVE1-depleted cells is the result of aberrant cell division. Considering that cultured *T. brucei* cells are asynchronous and that there are currently no established methods for synchronizing these cells that are compatible with RNAi, we developed a strategy to identify cells that had completed cell division prior to PAVE1 RNAi induction. We induced PAVE1 RNAi in log-phase cell cultures and harvested cells after 4 or 6 h of RNAi induction and restricted our analysis to 1N2K cells to select for cells that had progressed well into G1 phase at the time of PAVE1 depletion. The cell cycle for the 29.13 line carrying the PAVE1 RNAi hairpin is 12.9 h. Previous studies have shown that 1N2K cells would have initiated cell division approximately 10.8 h (0.84 units) prior to the time of harvest (38). Selecting 1N2K cells allowed us to eliminate any confounding effects on array morphology from cytokinesis and the posterior end remodeling that occurs in G1 phase immediately following cytokinesis, as these cells will have already completed these processes prior to PAVE1 RNAi induction (39,40). Thus, any defects observed in the array of 1N2K cells would be the direct result PAVE1 depletion without the combined effects of the major subpellicular array remodeling that occurs during the latest stages of cell division (Fig. 3C).

We visualized the array microtubules using immunofluorescence microscopy with an anti-α-tubulin antibody and measured the length of the array from the more posterior kinetoplast to the posterior edge of the cell in PAVE1 RNAi and control 1N2K cells. We found that the posterior array of PAVE1 RNAi cells at T=4 h were significantly shorter than T=0 h cells by an average of 0.99 ± 0.12 μm (Fig. 3D). Moreover, the average posterior array length of PAVE1 RNAi cells at T=6 h was 1.38 ± 0.23 μm shorter than T=0 h cells. However, the length of the array between the nucleus and posterior kinetoplast was the same at all three time points, which indicates that PAVE1 RNAi did not cause repositioning of the kinetoplast within the cell body or a shortening of the array at the cell midzone. These data suggest that PAVE1 is required to maintain the extended, tapering portion of microtubules at the cell posterior independent of its potential function during formation of the nascent cell posterior during cell division.

### PAVE1 immunoprecipitation reveals two potential interacting partners

The inter-microtubule crosslinking fibrils are approximately 6 nm in diameter and span the 18-20 nm distance between microtubules (18). We were curious if PAVE1 was part of a complex of proteins that formed the inter-microtubule crosslinks at the cell posterior. To determine if PAVE1 had interacting partners, we endogenously tagged both PAVE1 alleles with the fluorescent protein mNeonGreen (mNG) at their N-termini (Fig. 4A) and immunoprecipitated mNG-PAVE1 using mNeonGreenTrap antibody. Fractions of the eluate examined by SDS-PAGE and subsequent silver staining showed two primary unique bands compared to the control eluate (Fig. 4B). Mass spectrometry analysis of on-bead tryptic digests generated from mNG-PAVE1 and control immunoprecipitations revealed two potential interacting partners whose predicted molecular weight matched that of the unique bands on the silver-stained gel (Table 1). These potential interactors were an uncharacterized protein (Tb927.9.11540, predicted MW 51 kDa) and TbAIR9 (Tb927.11.17000, predicted MW 110 kDa).

**Figure 4.**
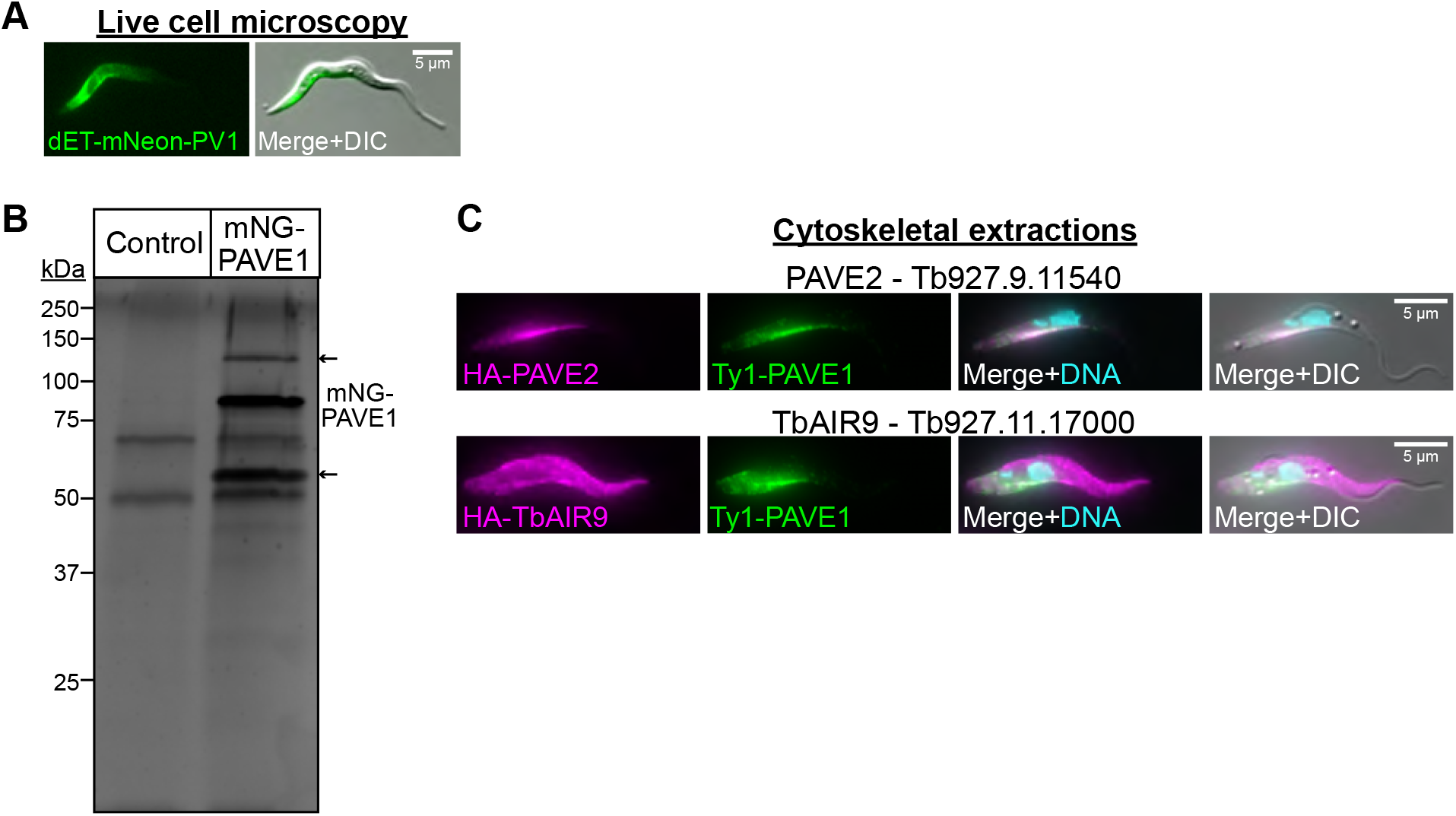
Immunoprecipitation of mNeonGreen-PAVE1 reveals two interacting partners. **(A)** Representative image of a live *T. brucei* cells expressing mNeonGreen-PAVE1 from both endogenous loci. **(B)** dET-mNeon-PAVE1 cells were harvested, lysed, and sonicated to solubilize cytoskeletal proteins. Clarified supernatant was batch-bound with magnetic camelid nanobodies against mNeonGreen. Beads were washed 6X and bound proteins were analyzed by mass spectrometry. Gel is a representative silver-stained SDS-PAGE gel of the elution from the mNeonGreen immunoprecipitation in control and mNeonGreen-PAVE1 samples. Both lanes are 50% of input material. Arrows indicate the two major unique protein bands present in the mNeonGreen-PAVE1 sample not present in WT 427 controls. **(C)** The two potential interacting partners of PAVE1 identified in the mNeonGreen-PAVE1 immunoprecipitation were endogenously tagged with the HA epitope tag. The cytoskeleton was extracted and cells were processed for immunofluorescence to co-localize each protein with Ty1-PAVE1.

We appended triple-HA epitope tags to the N-termini of the potential interactors in a cell line already harboring Ty1-tagged PAVE1, then performed cytoskeletal extractions followed by immunofluorescence microscopy. The uncharacterized HA-tagged protein Tb927.9.11540 was stably associated with the cytoskeleton and co-localized with Ty1-PAVE1, so we termed this new protein PAVE2 (Fig. 4C, top). HA-TbAIR9 was stably associated with the entire array, as previously reported (34). There was more intense staining of HA-TbAIR9 at the anterior portion of the array, where the Ty1-PAVE1 signal is minimal (Fig. 4C, bottom).

### PAVE1 and PAVE2 require each other for stability and localization

Like PAVE1, PAVE2 is conserved in kinetoplastids and has no homologs in other domains of life. PAVE2 was previously identified as a potential interactor of the cytokinetic regulator TOEFAZ1 in the same proximity-dependent biotinylation screen that originally identified PAVE1 (42). To test its function, we conducted RNAi against PAVE2 and found that cells were unable to divide after 4 d of RNAi induction (Supp. Fig. 2A & B). PAVE2 RNAi led to an accumulation of multinucleated cells (MultiN), cells with two nuclei and one kinetoplast (2N1K), and anucleate zoids (0N1K) (Fig. 5 A, B), which is similar to the cell division defect observed in PAVE1 RNAi (42).

**Figure 5.**
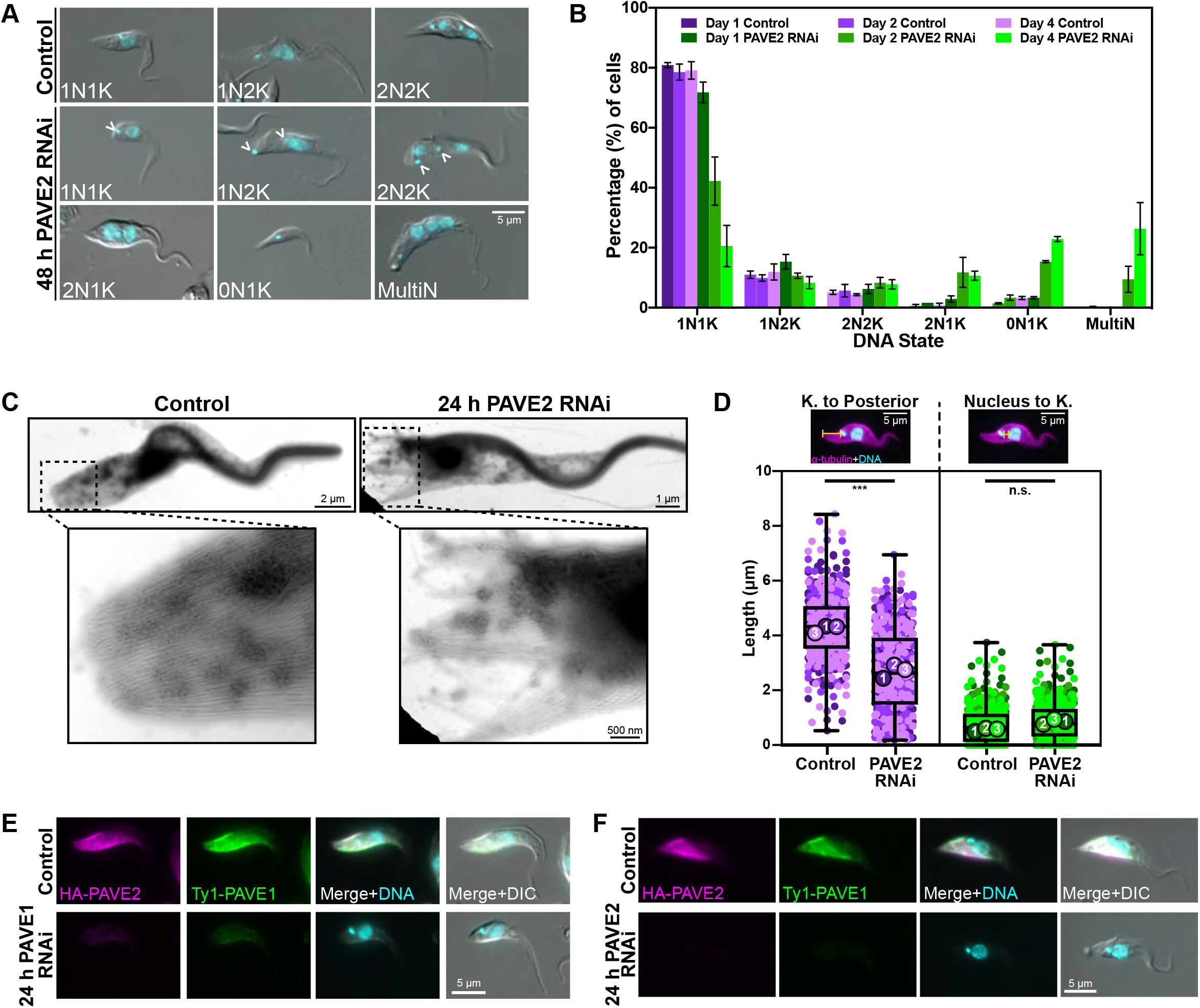
PAVE2 phenocopies PAVE1. **(A)** PAVE2 RNAi was induced for two days, after which control and PAVE2-depleted cells were fixed with paraformaldehyde and examined by epifluorescent microscopy. **(B)** Quantification of DNA states in control and PAVE2 RNAi cells after 1, 2, and 4 d of RNAi induction. Bars represent the mean and the error bars are the S.D. 300 cells in both control and PAVE2 RNAi conditions from N=3 independent biological replicates were counted at each time point. **(C)** Whole-mount transmission electron microscopy of control and PAVE2 RNAi cells after 1 d of PAVE2 depletion. **(D)** PAVE2 RNAi was induced for 1 d and cells were harvested, fixed in paraformaldehyde, and stained with anti-a-tubulin to demarcate the subpellicular array. Superplot is of the measured distance between the kinetoplast and the outer posterior edge of the array, and the posterior edge of the nucleus to the kinetoplast of 1N1K cells in control and PAVE2 RNAi conditions. 100 1N1K cells were measured for both control and PAVE2 RNAi conditions in N=3 independent biological replicates. Unpaired two-tailed Student’s *t-*tests were performed using the averages from N=3 independent biological replicates. ***P=0.0006. n.s. P=0.0753. **(E)** Control and PAVE1 RNAi cells were fixed in paraformaldehyde and stained after 1 d of PAVE1 depletion. PAVE2 does not localize to the array in the absence of PAVE1. **(F)** Control and PAVE2 RNAi cells were fixed in paraformaldehyde and stained after 1 d of PAVE2 depletion. PAVE1 does not localize to the array in the absence of PAVE2.

Strikingly, we saw that PAVE2 depletion resulted in cell bodies whose posterior ends directly abutted the kinetoplast (Fig. 5A, empty arrowheads). This mirrored the PAVE1 RNAi posterior truncation phenotype (42). The microtubules of truncated posterior arrays in PAVE2 RNAi cells were also disorganized in comparison to control cells when analyzed by TEM (Fig. 5C). We measured the length of the posterior array in control and PAVE2 RNAi cells to confirm the truncation phenotype. We found that the mean distance between the kinetoplast and posterior end of 1N1K cells was an average of 1.62 ± 0.31 μm shorter than controls after 24 h of RNAi induction, while the distance between the nucleus and kinetoplast remained constant (Fig. 5D). This suggested the posterior truncation phenotype was not due to aberrant kinetoplast placement or defects in the array at other locations but to the specific destabilization of microtubules at the posterior array, which phenocopies PAVE1 RNAi.

Both PAVE1 and PAVE2 RNAi result in the truncation of the posterior array, which suggests that these proteins may have a shared function or may form a complex. We depleted PAVE1 using RNAi and determined the localization of PAVE2 to establish if PAVE2 relied on PAVE1 for localization or stability within the cell. We found that HA-tagged PAVE2 localization to the posterior subpellicular array decreased as Ty1-PAVE1 protein levels decreased (Fig. 5E). Western blot analysis showed that PAVE2 protein levels also decreased during PAVE1 RNAi (Supp. Fig. 2C). We also found that the inverse relationship existed as well; when PAVE2 was depleted by RNAi, PAVE1 no longer localized to the posterior array (Fig. 5F) and its levels were reduced (Supp. Fig. 2D). These results suggest that PAVE1 and PAVE2 require each other for localization and protein stability inside the cell.

### PAVE1 and PAVE2 form a microtubule-associated complex *in vitro*

The interdependence of PAVE1 and PAVE2 *in vivo* (Fig. 5) and the near 1:1 stoichiometry of PAVE2 pulled down in the PAVE1 immunoprecipitation (Fig. 4B) suggest that the two proteins may form a protein complex. To test this, we co-expressed PAVE1 and PAVE2 recombinantly in *Escherichia coli*. The solubility-promoting maltose binding protein (MBP) followed by an oligo-His tag and TEV protease site were appended to the N-terminus of mNG-PAVE1, while a Strep peptide tag was added to the N-terminus of PAVE2. Lysates from *E. coli* co-expressors were incubated with nickel resin, followed by elution of the captured proteins and treatment with TEV protease to cleave the MBP-oligoHis tag from mNG-PAVE1. Subsequent capture of the eluate with Strep-Tactin resin resulted in a pure complex of mNG-PAVE1 and Strep-PAVE2, which we termed the PAVE complex (Fig. 6A, left). Attempts to purify either PAVE1 or PAVE2 on their own were not successful, which suggests that these proteins require each other for proper folding. The purified PAVE complex was diluted into a low salt buffer and then clarified by ultracentrifugation to test its solubility. The majority of the complex remained in solution, suggesting that the PAVE complex is highly soluble (Fig. 6A, right). These expression results, combined with their relationship in vivo, suggest that PAVE1 and PAVE2 form a hetero-oligomer that is responsible for the construction and maintenance of the tapered cell posterior.

**Figure 6.**
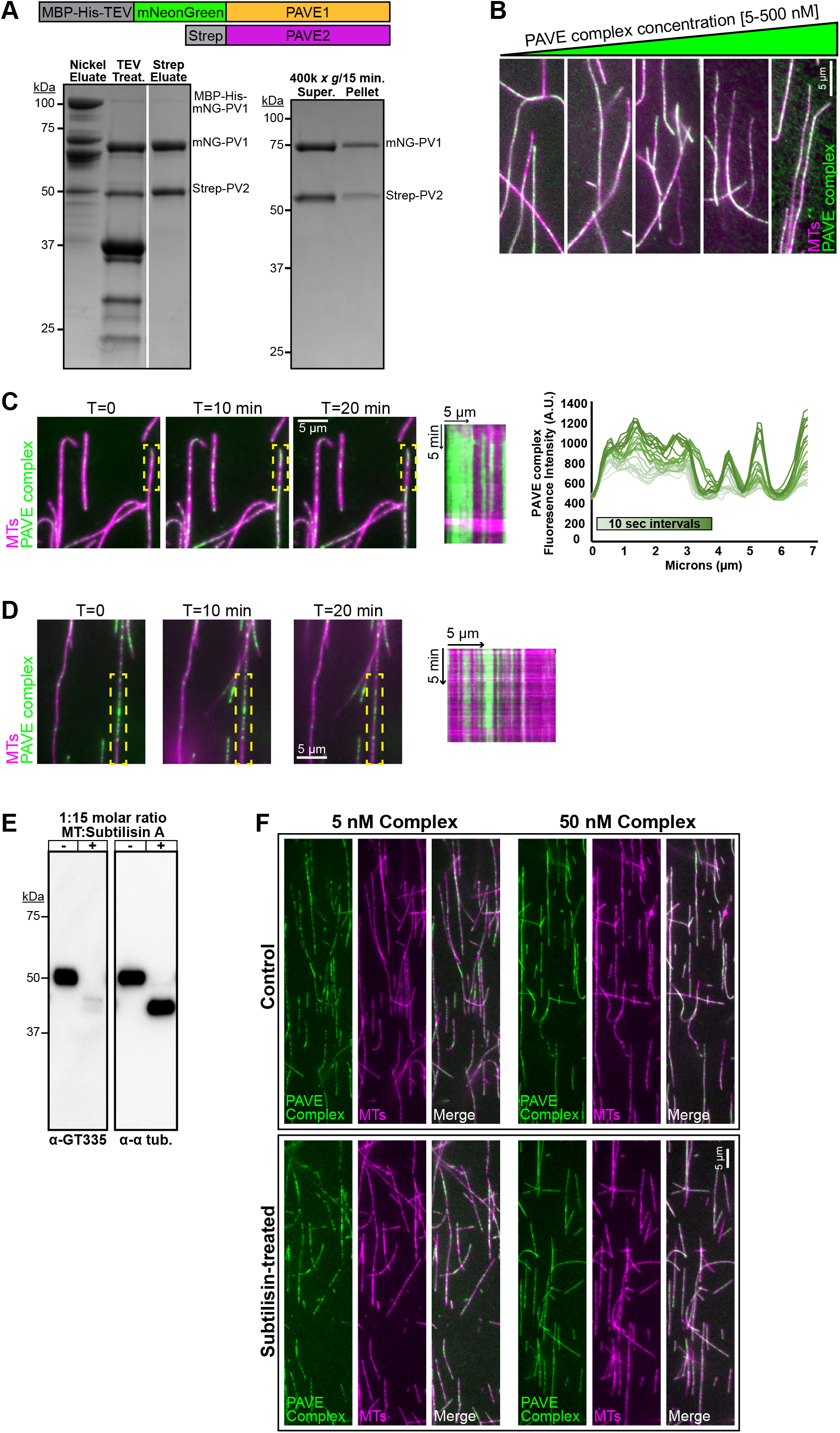
PAVE1 and PAVE2 form a complex in vitro that binds the microtubule lattice. **(A)** (Left) Coomassie-stained SDS-PAGE gel demonstrating the co-purification of mNeonGreen-PAVE1 and Strep-PAVE2 from *E. coli* cells. *E. coli* cells co-expressing [MBP-oligoHis-TEV]-nNeonGreen-PAVE1 and Strep-PAVE2 constructs were harvested, lysed, and clarified supernatant were batch-bound with Ni-NTA resin. The eluate was treated with TEV protease and allowed to interact with Strep-Tactin XT resin. Proteins were eluted using 50 mM biotin. The Strep eluate is 4 μg of protein. (Right) The PAVE complex was diluted into a low salt buffer containing 50 mM NaCl and ultracentrifuged. Equal fractions of both the supernatant and pellet were separated on an SDS-PAGE gel and stained with Coomassie blue. **(B)** 5, 50, 100, 200, and 500 nM clarified PAVE complex was allowed to interact with Taxol-stabilized bovine microtubules labeled with the Cy5 fluor for 20 min RT. The PAVE complex-microtubule solution was settled onto a blocked flow cell and unbound protein was washed away. The flow cell was imaged using epifluorescence microscopy. **(C)** (Left) A PEGylated flow cell was blocked and Taxol-stabilized bovine Cy5-microtubules were allowed to attach to the flow cell surface. 5 nM clarified PAVE complex was added to the flow cell chamber and immediately imaged using TIRF microscopy. Images were taken every 10 sec for 20 min. (Middle) Kymograph of PAVE complex on the microtubule outlined in yellow. (Right) Fluorescence intensity line scan of the microtubule outlined in yellow for the first 4 min 10 sec of TIRF imaging. **(D)** (Left) 5 nM clarified PAVE complex was mixed with Taxol-stabilized bovine Cy5-microtubules for 10 min RT to pre-seed patches. The PAVE complex-microtubule mixture was flowed into a blocked and PEGylated flow cell and allowed to attach to the flow cell surface. Unbound protein was washed away and the flow cell was immediately imaged using TIRF microscopy. Images were taken every 10 sec for 20 min. (Right) Kymograph of PAVE complex on the microtubule outlined in yellow. **(E)** Control microtubules and microtubules treated with subtilisin A were collected for western blotting. The post-translational modification of glutamylation is no longer present on subtilisin-treated microtubules, as detected with the antibody GT335, and there is a molecular weight shift showing that the C-terminal tails have been cleaved, as detected with anti-a-tubulin. **(F)** PAVE complex was incubated with control or subtilisin-treated Taxol-stabilized bovine Cy5-microtubules as in (B) and imaged using epifluorescence. There is no difference in PAVE complex binding to subtilisin-treated microtubules cleaved of their C-terminal tails in comparison to control, indicating that the PAVE complex binds directly to the microtubule lattice.

To determine if the PAVE complex can bind directly to microtubules, we incubated purified complex in solution with microtubules polymerized from bovine brain tubulin labeled with Cy5 dye (Fig. 6B). The PAVE complex-microtubule mixture was added to a flow cell and allowed to attach to the surface, after which unbound protein was washed away and the flow cell was imaged using epifluorescence microscopy. The PAVE complex bound to microtubules at concentrations as low as 5 nM (Fig. 6B). However, the PAVE complex did not crosslink microtubules into higher order structures such as bundles, regardless of the concentration of PAVE complex used or the length of the incubation. Although PAVE1 and thus PAVE2 as a direct binding partner localize to the inter-microtubule crosslinks of the *T. brucei* posterior subpellicular array in vivo, they do not appear to have the capacity to crosslink microtubules in vitro. This suggests that PAVE1 and PAVE2 are a necessary component of the inter-microtubule crosslinks at the posterior array but are not sufficient to form these crosslinks.

Although the PAVE complex cannot bundle microtubules, we observed that it localized to microtubules in patches even at high nanomolar concentrations, rather than being distributed in a uniform manner along the length of the microtubule (Fig. 6B). To determine how these patches form, we performed total internal reflection fluorescence (TIRF) microscopy to image the PAVE complex binding to microtubules. Cy5-labelled microtubules were attached to a flow cell, after which 5 nM PAVE complex was added into the chamber and immediately imaged using TIRF microscopy (Movie 1). We saw that the PAVE complex bound to microtubules in discrete patches; these patches did not grow over time, although they did increase in intensity (Fig. 6C, middle and right). Moreover, the PAVE complex patches were highly persistent. When patches were pre-seeded on microtubules in solution, added to a flow cell, and unbound complex was washed away, the pre-seeded patches remained for greater than 20 min (Fig. 6D, Movie 2). These results demonstrate that the PAVE complex binds to microtubules in static patches, which suggests that the complex may recognize specific regions of microtubules.

Microtubules have many post-translational modifications (PTMs), which can differentially recruit MAPs or affect MAP behavior (26). Most microtubule PTMs occur on their highly disordered and charged C-terminal tails. We cleaved the C-terminal tails from bovine Cy5-labelled microtubules using subtilisin protease (46) to establish if the PAVE complex recognized tail-associated PTMs, which could explain its patchy localization pattern. We confirmed the removal of tubulin C-terminal tails by subtilisin using western blotting analysis (Fig. 6E). We found that glutamylation, which is a modification that occurs exclusively on the C-terminal tail of α- and β-tubulin, was significantly reduced by subtilisin treatment. Blotting with anti-tubulin antibody also showed that the protease-treated tubulin had undergone a molecular weight shift, indicating that the C-terminal tail had been removed. We incubated the PAVE complex with control and subtilisin-treated microtubules and found no difference in the intensity or pattern of binding, regardless of the concentration of PAVE complex used (Fig. 6F). These results suggest that the PAVE complex binds directly to the microtubule lattice and may recognize a specific lattice structure rather than a post-translational modification in the tubulin tails.

### TbAIR9 controls the distribution of PAVE1 in the subpellicular array

TbAIR9, which localizes to the entire subpellicular array, was also identified as a potential PAVE1 interactor in our immunoprecipitation analysis (Fig. 4C, bottom). TbAIR9 shares sequence similarity with the microtubule associated protein AIR9 found in *Arabidopsis thaliana*, which has a role in positioning the phragmoplast during plant cytokinesis (47). We tested whether depleting TbAIR9 had an effect on PAVE1. We conducted RNAi against TbAIR9 in a cell line where TbAIR9 was tagged with HA and PAVE1 was tagged with Ty1. Cells lacking TbAIR9 developed aberrant cell morphologies and ceased to divide, as previously reported (Supp. Fig. 3A & B) (34). After 24 h of RNAi induction, HA-tagged TbAIR9 signal remained at the cell anterior while the cell posterior lacked the protein (Supp. Fig. 3C). In uninduced control cells, Ty1-tagged PAVE1 strongly localized to the posterior and ventral edge of the cell, while in the absence of TbAIR9, PAVE1 lost its preference for this subdomain and spread throughout the entirety of the array (Fig. 7A). A line scan measuring PAVE1 intensity placed along the ventral edge of TbAIR9 RNAi cells showed that PAVE1 labeling was less intense at the cell posterior and more intense at the cell anterior in comparison to control cells (Fig. 7B). PAVE1 protein levels were not altered, indicating that TbAIR9 depletion only affected the ability of PAVE1 to localize to the posterior subdomain (Fig. 7C). Depleting PAVE1 from cells did not affect TbAIR9 stability or localization, except for its absence at the posterior subpellicular array in cells where the posterior was truncated (Supp. Fig. 3D & E).

**Figure 7.**
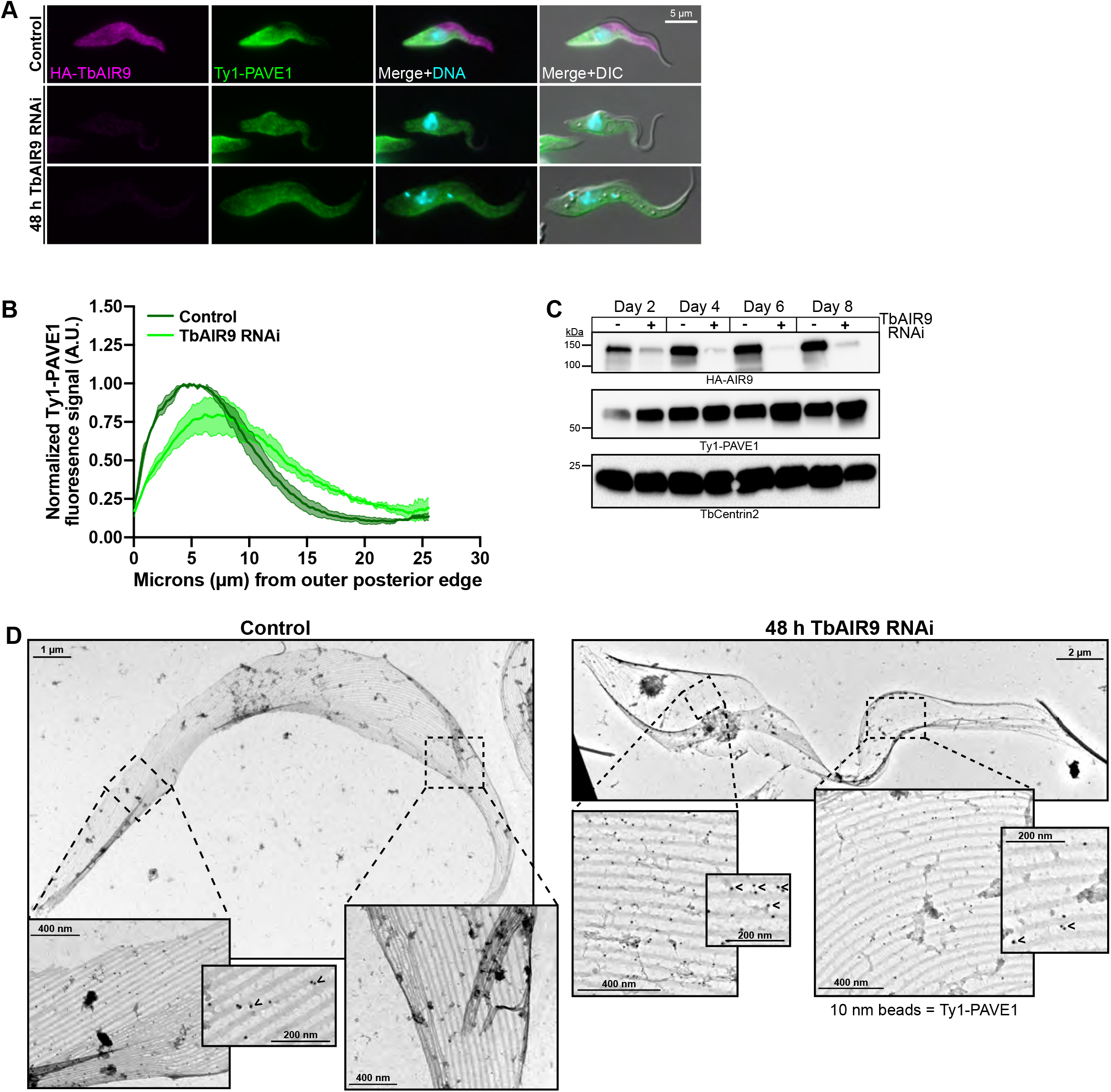
TbAIR9 confines PAVE1 to the crosslinks at the cell posterior. **(A)** Control and TbAIR9 RNAi cells were fixed in paraformaldehyde and stained after 2 d of TbAIR9 depletion. Ty1-PAVE1 signal appears spread throughout the array in the absence of TbAIR9. **(B)** The fluorescence signal intensity of Ty1-PAVE1 was measured along the ventral side of control and TbAIR9 RNAi cells after 2 d of TbAIR9 depletion. The signal intensities were normalized to the maximum control intensity, averaged, and plotted with the S.D. 50 1N1K cells in both control and TbAIR9 RNAi conditions from N=3 independent biological replicates were measured. Ty1-PAVE1 signal is more evenly distributed throughout the array in TbAIR9 depleted cells than in control cells. **(C)** TbAIR9 RNAi was induced and lysates were collected every 48 h from control and TbAIR9 depleted cells. Lysates were separated by SDS-PAGE and transferred to nitrocellulose for western blotting. Ty1-PAVE1 signal remains stable during TbAIR9 depletion. **(D)** Subpellicular array sheets of control and TbAIR9 RNAi cells after 2 d of TbAIR9 depletion immunogold labeled against Ty1-PAVE1 with 10 nm beads. Ty1-PAVE1 localizes to the inter-microtubule crosslinks at both the posterior and anterior ends of the array in the absence of TbAIR9.

The mis-localization of PAVE1 could be due to the disruption of inter-microtubule crosslinks during TbAIR9 depletion. To determine if crosslinks are still formed in cells depleted of TbAIR9, we created subpellicular array sheets of uninduced control and TbAIR9 RNAi cytoskeletons immunogold-labeled for Ty1-PAVE1 (Fig. 7D). In control cells, crosslinks were present throughout the array and PAVE1 localized to those present at the cell posterior (Fig. 7D, left). In TbAIR9 RNAi cells, crosslinks were also still present in the entire array (Fig. 7D, right). However, PAVE1 now localized to the crosslinks at both the posterior and anterior ends of the cell. This result suggests that TbAIR9 is not necessary to build the inter-microtubule crosslinks present in the array but is required to properly limit the distribution of PAVE1 to the cell posterior.

### TbAIR9 is a global regulator of subpellicular array-associated protein localization

The striking redistribution of PAVE1 throughout the array during TbAIR9 depletion led us to consider that TbAIR9 may regulate the localization of other cytoskeletal proteins in the subpellicular array. To test this, we identified two proteins that localized to different domains of the array using the whole-genome localization database TrypTag (48).

Tb927.9.10790 is a 33 kDa protein annotated by TrypTag as localizing to the middle of the subpellicular array. This protein is found only within the genomes of *T. brucei* and *T. cruzi* with no homologs present in other domains of life. Tb927.11.1840 is a 45 kDa protein that localizes to the anterior array in TrypTag and appears to be unique to kinetoplastids (49). We endogenously tagged both proteins at their C-termini and performed immunofluorescence on extracted cytoskeletons. Both 10790-Ty1 and 1840-Ty1 were stably associated with the cytoskeleton and localized to the midzone and anterior domains of the array, respectively (Fig. 8A).

**Figure 8.**
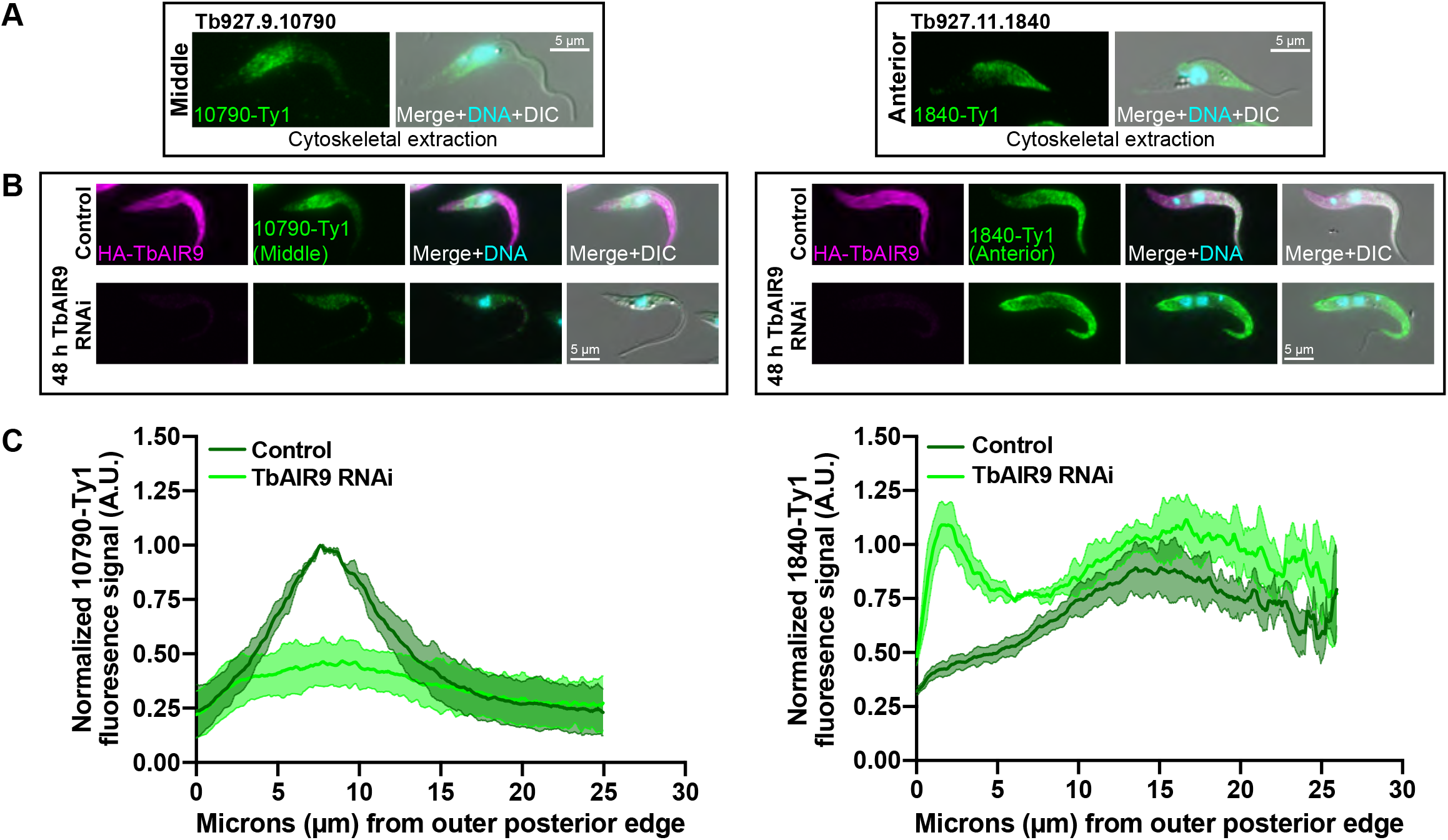
TbAIR9 maintains the localization array-associated proteins at specific domains. **(A)** Identification of array-associated proteins from TrypTag that localize to the middle (Tb927.9.10790, left) and anterior (Tb927.11.1840, right) domains of the subpellicular array. Both proteins were tagged at their C-terminus with Ty1 and cytoskeletal extractions followed by immunofluorescence with anti-Ty1 antibody was perfomed. Both proteins remained bound to the array in cytoskeletal extractions. **(B)** Cell lines containing 10790-Ty1 (left) and 1840-Ty1 (right) endogenous tags were fixed in paraformaldehyde and stained after 2 d of TbAIR9 depletion. 10790-Ty1 signal is depleted and spread throughout the cell body in the absence of TbAIR9, while 1840-Ty1 signal is detected at the posterior and anterior ends in TbAIR9 depleted cells. **(C)** Quantification of 10790-Ty1 (left) and 1840-Ty1 (right) fluorescent signal intensity along the ventral side of the cell body in TbAIR9 RNAi cells after 2 d of TbAIR9 depletion. The signal intensities were normalized to the maximum control values and plotted with the S.D. 50 1N1K cells in both control and TbAIR9 RNAi conditions from N=3 independent biological replicates were measured.

We next determined if TbAIR9 is necessary to confine the distribution of 10790 and 1840 to their respective domains of the subpellicular array, as it does in confining PAVE1 to the array posterior (Fig. 7). We induced TbAIR9 RNAi and localized 10790 and 1840 using immunofluorescence microscopy. We found that upon TbAIR9 depletion, 10790 is weakly distributed throughout the array (Fig. 9B & C, left), while 1840 localized to both the posterior and anterior array, with weaker signal at the array midzone (Fig. 8B & C, right). TbAIR9 RNAi decreased the protein levels of 10790 (Supp. Fig. 4A), while 1840 protein levels increased (Supp. Fig. 4B). These results suggest that TbAIR9 functions to organize and confine array-associated proteins to distinct domains within the array.

## DISCUSSION

In this work, we have characterized a series of proteins that appear to localize to and regulate the inter-microtubule crosslinking fibrils of the subpellicular array, which previously had no known components. These results show that the subpellicular array is organized into subdomains that are likely defined by microtubule crosslinks with unique protein compositions, which may differentially structure the array to shape *T. brucei* cells and alter the local properties of the microtubules. Asymmetric protein localization within the basal bodies of *Tetrahymena* stabilizes them against ciliary beating forces (50) and is required for basal body replication and positioning in *Paramecium* (51,52). *T. brucei* may employ a similar mechanism on a broader scale across the array to withstand the forces from flagellar beating and maintain array shape.

Our quantitative iEM shows that PAVE1 preferentially localizes to the ventral side of the subpellicular microtubules and the inter-microtubule crosslinks at the cell posterior (Fig. 2C-E). While our ‘unroofing’ approach disrupts many of the inter-microtubule crosslinking fibrils, the spacing and length of the structures we observe coincide with the placement of the inter-microtubule crosslinks found by others in subpellicular array sheets (16,53). Currently, there are no methods available for immuno-labeling intact inter-microtubule crosslinks, which are best visualized using electron tomography or quick-freeze deep-etch EM (7,18,54). The specific localization of PAVE1 to the ventral side of microtubules suggests that the crosslinking fibrils are composed of asymmetrically distributed proteins and that PAVE1 contacts the microtubule on one side of the crosslink. This placement argues that there must be other proteins that span the inter-microtubule distance and contact the dorsal side of the adjacent microtubule to complete the crosslink. This is in line with our in vitro data, which show that the PAVE complex is not able crosslink microtubules into bundles in solution, most probably because the complex cannot assemble a complete span that can contact two microtubules simultaneously (Fig. 6B). Instead, our data suggest that the PAVE complex is an essential component of the ventral microtubule interface of the crosslinks at the cell posterior. The asymmetric composition of array crosslinks has also been suggested by the array-associated protein WCB, which preferentially localizes to one side of the plasma-membrane facing surface of array microtubules (35,53). Additional proteins that can contribute to the assembly of a complete crosslink are currently being sought.

PAVE1 is initially loaded onto the subpellicular microtubules at the extreme posterior end of the cell (Fig. 3A). The growing plus-ends of subpellicular array microtubules are concentrated at this location throughout the cell cycle(16,39). The microtubule plus-ends are recruitment sites for new tubulin dimers, which are preferentially loaded with GTP and still contain the terminal tyrosine residue on their C-terminal tails (43). Removing the C-terminal tails with subtilisin had no effect on the avidity or pattern of PAVE complex binding to microtubules, which argues that the complex binds directly to the microtubule lattice (Fig. 6F). It is intriguing that the PAVE complex binds to microtubules in patches, even at high concentrations of protein (Fig. 6B). The PAVE complex is unlikely to be a microtubule plus-end binding protein, as its localization is not exclusive microtubules that still contain terminal tyrosination (Fig. 1A). Recently, it has been suggested that several MAPs, including the plus-end binder EB1 and the neuronal MAP doublecortin, preferentially recognize more open and irregular conformations of the microtubule lattice, which are often present at microtubule plus-ends and sites of curvature or damage (27,28,55). The neuronal MAP tau also forms concentration-dependent condensates on the GDP-lattice of microtubules, especially at sites of curvature, which protects the microtubule from the action of severing enzymes (56,57). It is possible that PAVE1 may recognize irregularities in the microtubule lattice, which are concentrated at the polymerizing end of the microtubule where the lattice is incomplete. Finally, it is also possible that polymerizing microtubule ends are the only places with free microtubule binding sites in the subpellicular array, as sites located more anterior in the array are already occupied with different MAPs.

Our HaloTag capping assay shows that the inter-microtubule crosslinks are not static within the array (Fig. 3A). PAVE1 migration towards the anterior end suggests two possible mechanisms of inter-microtubule crosslink movement. One possibility is that the inter-microtubule crosslinks remain associated with specific parts of the microtubule lattice and that the microtubules turn over as new tubulin is added to the cell posterior, which moves the crosslinks toward the anterior end of the cell. The other possibility is that inter-microtubule crosslinks can diffuse along the microtubules and that the selective recruitment of new PAVE1 to the extreme posterior end drives the movement of extant crosslinks specifically towards the cell anterior. Our in vitro results suggest that the PAVE complex forms persistent contacts with microtubules and does not diffuse along their lengths (Fig 6C &D). This, along with the fact that new tubulin subunits are constantly added to the cell posterior, argues that PAVE1 is forming persistent contacts with the microtubules that anchor the crosslinks in place and suggests the former possibility of crosslink movement. The rapid rate of posterior end retraction upon PAVE1 depletion (Fig. 3C) argues that new inter-microtubule crosslinks are constantly being assembled and are essential for retaining the extended shape of the cell. Since the subpellicular microtubules do not appear to undergo catastrophes that shorten them, this observation requires that cells that are not undergoing cell division have some process that truncates or removes tubulin subunits from the array at a rate similar to that which tubulin is added to the posterior end. Recently, it has been theorized that the microtubules of the array may ‘slide’ to the anterior end of the cell to accommodate remodeling events after cell division, which could be suggestive of a microtubule treadmilling process in the array (40).

Since PAVE1 does not localize to the anterior end of the cell, some mechanism must exist to remove it from the microtubule crosslinks once the crosslinks reach the cell midzone. It is currently unknown if the crosslinks are remodeled by replacing subdomain-specific proteins such as PAVE1/2 with other proteins, or if new crosslinks are constructed at each subdomain, which would require a mechanism for installing them within the local array subdomain. It would also require that each subdomain of the subpellicular array have specialized sites for the local construction of different microtubule crosslinks, which have not been observed yet.

TbAIR9 is associated with the PAVE complex, which argues that the protein is also a component of the inter-microtubule crosslinks (Figure 4B). The trypanosomatid homolog of AIR9 differs from its plant counterpart in that it lacks a microtubule binding domain at its N-terminus and has fewer A9 repeats at its C-terminus, which are Ig-like domains that likely serve as sites for protein-protein interactions (58). The absence of the microtubule binding domain suggests that TbAIR9 is recruited to the microtubules by other proteins, such as the PAVE complex, likely via the A9 repeats.

The broad distribution of TbAIR9 across the array (Fig. 4C) suggests that it may be a universal component that makes up a core part of the microtubule crosslinking fibrils, whose behavior are then locally tuned by proteins that interact with TbAIR9. While the PAVE complex does not appear to bind to post-translational modifications on the C-terminal tail of tubulin, other subpellicular components may use them to localize to different regions of the array and then form associations with core crosslink-associated proteins such as TbAIR9. Tubulin post-translational modifications such as polyglutamylation and tyrosination have different distribution patterns along the array and may serve as landmarks for recruiting specific MAPs (16,39,59,60).

In *Arabidopsis*, AtAIR9 localizes to the cortical microtubules, which are analogous to the subpellicular array in that they underlie the plasma membrane. Cortical array microtubules in plants primarily control the distribution of cellulose synthetase and possibly act as tracks for membrane traffic (61). Deletion of AtAIR9 does not produce a phenotype, but a strong synthetic effect is observed with the simultaneous deletion of a second microtubule-binding protein known as TANGLED1, which produces plants with disorganized cortical microtubules and twisted roots (62). Depletion of TbAIR9 did not cause the crosslinks to disappear (Fig. 7D), so TbAIR9 function must be focused on controlling the distribution of proteins along the array (Fig. 8). The phenotype of TbAIR9 depletion is complex, producing anucleate and multinucleated cells with elongated and epimastigote-like morphologies (Supp. Fig. 3B) (34). Our data show that TbAIR9 depletion results in the mislocalization PAVE1 (Fig.7) and two newly identified array-associated proteins, Tb927.9.10790 and Tb927.11.1840 (Fig.8). Each of these proteins is confined to different subdomains of the array in wild-type cells, but the subdomain architecture is lost or scrambled in the absence of TbAIR9. This suggests that the TbAIR9 RNAi phenotype is the result of the inability to locally regulate microtubule properties in these subdomains, which appears to have a strong effect on parasite cell division.

Our work has highlighted the potential functions of a set of proteins that are components of the inter-microtubule crosslinking fibrils within the subpellicular array. These crosslinks appear consist of different protein components that likely have discrete functions in order to locally tune array dynamics. While the PAVE complex appears to function to maintain the length and shape of the array microtubules at the cell posterior, the functions of Tb927.9.10790 and Tb927.11.1840 are yet to determined. Future work is also needed to discover how TbAIR9 organizes these proteins into defined domains across the array. The TrypTag database contains many previously unidentified proteins that appear to localize to the subpellicular array; some are present along the whole array, and others localize to subdomains. It is interesting to note that the distribution patterns of some of these proteins are more complex, including more variety along the dorsal and ventral sides. Uncovering the function of these proteins is essential to understanding subpellicular array biogenesis and how this structure maintains its shape throughout the cell cycle and during the cell body waveforms created by *T. brucei* motility.

## MATERIALS AND METHODS

### Antibodies

Antibodies are from the following sources and were used at the following dilutions: anti-Ty1 (1:1000) from Sebastian Lourido (Massachusetts Institute of Technology—Boston, MA), TAT1 (1:100) from Jack Sunter (Oxford Brookes University—Oxford, United Kingdom). YL1/2 (1:4,000) was purchased from ThermoFisher Scientific (Waltham, MA, cat# MA1-80017) and anti-a tubulin (1:10,000) was also purchased from ThermoFisher Scientific (clone B-5-1-2).

Anti-HA antibody (1:500) was purchased from Sigma-Aldrich (St. Louis, MA, clone 3F10). GT335 (1:25,000) was purchased from Adipogen (San Diego, CA, cat# AG-20B-0020-C100). Anti-TbCentrin2 (1:150) antibody was previously described (63).

### Cell Culture

Wild-type procyclic *T. brucei brucei* 427 strain cells and the doxycycline-inducible 29-13 *T. brucei* cell line (64) were used to perform experiments. 427 cells were passaged in Beck’s Medium (Hyclone—GE Healthcare, Logan, Utah) supplemented with 10% fetal calf serum (Gemini Bioproducts—West Sacramento, CA), while 29-13 cells were passaged in Beck’s Medium supplemented with 15% doxycycline-free fetal calf serum (R&D Systems—Minneapolis, MN), 50 μg mL^-1^ hygromycin (ThermoFisher Scientific), and 15 μg mL^-1^ G418 (Sigma-Aldrich). 427 and 29-13 media also included 10 μg mL^-1^ gentamycin (ThermoFisher Scientific) and 500 μg mL^-1^ penicillin-streptomycin-glutamine (ThermoFisher Scientific). All cells were maintained at 27 °C. Cells were counted using a particle counter (Z2 Coulter Counter, Beckman Coulter—Brea, CA).

### Cloning and Cell Line Construction

All constructs were created by PCR amplification of inserts from *T. brucei* genomic DNA using Q5 polymerase (NEB—Ipswich, MA) followed by either restriction-ligation or Gibson Assembly into a sequencing vector (PCR4Blunt) or a modified pLEW100 for long-hairpin RNAi (lhRNAi) generation as previously described (65). Each DNA construct was validated by sequencing and transfected into cells using an electroporator (GenePulser xCell, Bio-Rad— Hercules, CA). Clonal cell lines were created by selection and limiting dilution. Cell lines were validated with western blotting and loci PCRs. All sequences were obtained from TriTrypDB (49).

#### RNAi constructs

Sequences for RNAi were selected from the coding sequence of the target protein using the RNAit online tool (https://dag.compbio.dundee.ac.uk/RNAit/) (66). 400-600 base pairs were used to create lhRNAi constructs as follows: PAVE1 (Tb927.8.2030)—bp 97-543; PAVE2 (Tb927.9.11540)—bp 514-1112; TbAIR9 (Tb927.11.17000)—bp 1868-2303. Constructs were linearized for transfection with NotI (NEB) and transfected into the 29-13 cell line, after which clonal populations were obtained by selection with 40 μg ml^-1^ Zeocin (Life Technologies— Carlsbad, CA).

#### Endogenous tagging constructs

Endogenous tagging constructs were created using Gibson Assembly into the PCR4Blunt sequencing vector. N-terminal tagging constructs (PAVE1, PAVE2, TbAIR9) were targeted to their respective endogenous loci using last 500 bp of 5’ UTR and first 500 bp of the coding sequence. C-terminal tagging constructs (Tb927.9.10790, Tb927.11.1840) were targeted to their loci using the last 500 bp of the coding sequence followed by the first 500 bp of the 3’ UTR. Endogenous tagging constructs were excised using PacI and NsiI (NEB) and transfected into either the 427 or 29-13 cell lines, and clonal populations were selected using 10 μg mL^-1^ blasticidin or 1 μg mL^-1^ puromycin.

### Image Acquisition for Fluorescence Microscopy

#### Epifluorescence acquisition

Images were acquired using a Zeiss Axio Observer.Z1 microscope (Carl Zeiss Microscopy—Oberkochen, Germany) using an 100X/1.4 NA Plan-Apochromat oil lens with an ORCA-Flash 4.0 V2 CMOS camera (Hamamatsu—Shizuoka, Japan). *Z*-stacks were acquired for each image.

#### Total Internal Reflection Fluorescence acquisition

Images were acquired at RT with the same Zeiss Axio Observer.Z1 microscope described above, equipped with a spinning illumination ring VectorTIRF system which includes a 405/488/561/640 quad band dichroic filter (Intelligent Imaging Innovations, Inc.—Denver, CO) and an Alpha Plan-Apochromat 100X/1.46 NA oil TIRF objective. Samples were excited using 488 nm 150mW and 647 nm 120mW lasers at 20% power. Images were acquired every 10 sec for 20 min using a Prime 95B Back Illuminated Scientific CMOS camera (Teledyne Photometrics—Tuscon, AZ) and a Definite Focus.2 system (Carl Zeiss Microscopy) for automatic focus correction.

SlideBook 6 digital microscopy software (Intelligent Imaging Innovations, Inc.) was used to manipulate the microscope and acquire images. Images were analyzed with ImageJ (National Institutes of Health—Bethesda, MD) and exported to Adobe Photoshop and Illustrator (CC 2020) for publication.

### Immunofluorescence Microscopy

Cells were harvested from culture by centrifugation at 2400 *x g* for 5 min and washed with PBS. Cells were centrifuged onto coverslips and fixed as per the conditions below. All fixation conditions for each experiment are specified in the text and figure legends. After fixation, cells were washed 3X with PBS and blocked using 5% goat serum (ThermoFisher) in PBS. Coverslips were then incubated with primary antibody diluted into blocking buffer for 1 hr RT, after which they were washed 3X with PBS and incubated with secondary antibody conjugated to Alexa Fluor −647, −568, and −488 (Life Technologies) diluted into blocking buffer for 1 hr RT. Coverslips were washed for a final 3X with PBS and mounted for epifluorescence imaging using Fluoromount-G with DAPI (Southern Biotech—Birmingham AL).

#### Methanol fixation

After cells were centrifuged onto coverslips, excess PBS was wicked away with tissue paper and the coverslips were quickly plunged into ice-cold methanol at 20 °C for 20 min.

#### Paraformaldehyde fixation

Coverslips were inverted onto a drop of 4% paraformaldehyde in PBS for 20 min RT in a humidified chamber. Cells were permeabilized by moving the coverslip to a drop of 0.25% NP-40 (IPEGAL CA-630, Sigma Aldrich) in PBS for 5 min RT.

#### Cytoskeletal extraction

Coverslips were inverted onto a drop of cytoskeletal extraction buffer (0.1 M PIPES pH 6.9, 2 mM EGTA, 1 mM MgSO4, 0.1 mM EDTA, 0.25% NP-40) for 2 min RT. Coverslips were then quickly washed 6X with PBS and fixed for 20 min RT with 4% paraformaldehyde in PBS.

### Transmission Electron Microscopy

#### Whole-mount extracted cytoskeletons preparation

Cells were harvested by centrifugation at 800 *x g* and washed 3X with PBS. Cells were briefly spun onto glow-discharged formvar- and carbon-coated grids (Electron Microscopy Sciences—Hatchfield, PA) at 800 *x g*. Grids were floated on two subsequent drops of cytoskeletal extraction buffer (0.1 M PIPES pH 6.9, 2 mM EGTA, 1 mM MgSO4, 0.1 mM EDTA, 1% NP-40) for 3 min RT each drop, and then briefly washed in a drop of extraction buffer without detergent. Grids were fixed in a drop of 2.5% glutaraldehyde in PBS for 5 min RT, after which they were washed with distilled water and negatively stained with 1% aurothioglucose (Sigma Aldrich).

#### Immunogold labeling

Whole-mount extracted cytoskeletons were prepared as above, but prior to fixation, cells were blocked by quickly moving the grids through 5 drops of blocking buffer (2% bovine serum albumin in PBS). Grids were incubated for 1 hr RT in primary antibody diluted in blocking buffer, after which they were washed 5X with blocking buffer. Grids were then incubated with 10 nm or 20 nm gold-conjugated secondary antibodies (BBI Solutions—Newport, United Kingdom) diluted in blocking buffer for 1 h RT. Grids were fixed and stained as above.

#### Subpellicular sheeting

Cells were prepared for sheeting as described in **(16)**. After cells were spun onto grids and extracted as above, the grids were gently fixed in a primary fixative of 3.2% paraformaldehyde in cellular extraction buffer without detergent for 5 min RT. The paraformaldehyde was then quenched by incubating the grids in 20 mM glycine in PBS for 5 min RT. Grids were blocked and immunogold labeled as above. After labeling, grids were fixed with 2.5% glutaraldehyde in PBS for 5 min RT, washed 2X with distilled water, and positively stained in a drop of 2% aqueous uranyl acetate for 45 min RT. Grids were dehydrated in an ethanol series and critical point dried. The top layer of the subpellicular array was removed to ‘unroof’ the cells by inverting the grid onto double-sided tape, applying gentle pressure, and then lifting the grid off the tape.

#### Image acquisition for TEM

Images were taken on a Phillips 410 transmission electron microscope at 100 kV equipped with a 1k x 1k Advantage HR CCD camera from Advanced Microscopy Techniques (AMT) using AMT imaging software. Images were analyzed in ImageJ and exported to Adobe Photoshop and Illustrator for processing.

#### Quantification of immunogold label distribution on subpellicular sheets

Micrographs of subpellicular sheets in which the flagellum was still attached were analyzed in ImageJ. The sheets were oriented along the dorsal-ventral axis using the flagellum as a fiducial marker. A 1 μm^2^ area of the sheet was selected for analysis. Each microtubule within the 1 μm^2^ area was traced using the segmented line tool. A semi-automated macro was written in ImageJ which converted the segmented line into a spline-fit line and straightened the microtubule according to the spline curve. A rectangle was placed on the bounding edges of the straightened microtubule and its centroid was calculated and plotted. The location of each gold bead on the microtubule was manually counted as dorsal (above the centroid line), centroid (touching the centroid line) or ventral (below the centroid line), as well whether the gold bead localized to an inter-microtubule crosslink. Three subpellicular array sheets were analyzed per N=3 independent biological replicates for each experiment. The data were recorded in Microsoft Excel and statistical tests were performed and the data graphed using Prism 8 (Graphpad).

### HaloTag Capping Assay

HaloTag **(45)** substrate conjugated to JF646 (Promega—Madison, WI) was added to a culture of 2 x 10^6^ cell mL^-1^ log-phase HaloTag-PAVE1 cells at a final concentration of 2 μM for 1 h at 27 °C. Cells were collected by centrifugation at 2400 *x g* and washed 3X with Beck’s Medium. Cells were resuspended at 2 x 10^6^ cell mL^-1^ in complete media and HaloTag substrate conjugated to TMR (Promega) was added at a final concentration of 5 μM. Cells were harvested at T=0 h, T=1 h, T=3 h, and T=8 h of TMR substrate addition by centrifugation at 2400 *x g*. Cells were washed 3X with HBSS, centrifuged onto coverslips, and fixed for 20 min RT with 4% paraformaldehyde in PBS. Coverslips were washed 3X with PBS and mounted with Fluoromount-G containing DAPI.

#### Image acquisition and fluorescence intensity analysis

Images were captured using epifluorescence as above. Maximum projections of each *Z*-stack were created and background-corrected using rolling ball background subtraction (radius = 100 pixels). The ventral edge of 50 1N1K cells per time point was traced using the segmented line tool. The fluorescence intensity along each line for HaloTag substrates conjugated to JF646 and TMR were recorded and exported to Microsoft Excel. The average pixel intensity and S.D. at each position along the line was calculated and normalized to 1 from N=3 independent biological replicates and graphed using Prism 8 software.

### RNAi

Log-phase cultures of 29-13 cells containing lhRNAi constructs were seeded at 1 x 10^6^ cell mL^-1^. RNAi was induced in using 1 μg mL^-1^ doxycyline (ThermoFisher Scientific) or 70% ethanol as a vehicle control. Cell growth was recorded every 24 h unless otherwise noted, and samples were collected for western blotting or immunofluorescence. RNAi cultures were re-seeded every 48 h in fresh media and doxycycline or ethanol was added as above. All RNAi generation plots represent three independent biological replicates and the error bars are the S.D.

#### PAVE1 and PAVE2 RNAi image analysis

Control and RNAi cells were fixed in methanol or paraformaldehyde and stained as above. Maximum projections of each *Z*-stack were created for analysis. For the PAVE1 RNAi short time course: cells were harvested and fixed in methanol after T=0 h, 4 h, and 6 h after RNAi induction. Only 1N2K cells were selected for analysis, as these cell types have progressed over halfway through the cell cycle at the time of harvest **(67)**, and defects in the array would be due solely to PAVE1 RNAi, and not defects in cytokinesis or G1 subpellicular array remodeling. 50 1N2K cells per each time point in N=3 independent biological replicates were measured. The distance from the posterior kinetoplast to the outer posterior edge of the subpellicular array, as well as the posterior nucleus edge to the posterior kinetoplast, were measured. For PAVE2 RNAi: Cells were harvested and fixed in paraformaldehyde. For DNA state analysis, 300 cells for each time point in control and PAVE2 RNAi conditions were counted in N=3 independent biological replicates. For posterior end measurements, 100 1N1K cells in control and Day 1 PAVE2 RNAi conditions were measured as above in N=3 independent biological replicates. Data were recorded in Microsoft Excel and statistical tests were conducted and the data graphed using Prism 8.

#### PAVE1, Tb927.9.10790, Tb927.11.1840 fluorescence intensity analysis during TbAIR9 RNAi

Control and TbAIR9 RNAi cells were harvested after two days of RNAi induction and fixed in paraformaldehyde and stained as above. The ventral side of 1N1K cells was traced using the segmented line tool, and fluorescence intensities were recorded and analyzed as above and normalized to the maximum control value. 50 1N1K cells each were measured in control and TbAIR9 RNAi conditions for N=3 independent biological replicates.

### SDS-PAGE and Western Blotting

*T. brucei* cells were collected by centrifugation at 2400 *x g*, washed with PBS, lysed in SDS-PAGE lysis buffer and incubated for 10 min at 99 °C. 3 x 10^6^ cell equivalents/lane were separated by SDS-PAGE and transferred to a nitrocellulose membrane. Blots were blocked with 5% (w/v) non-fat dry milk dissolved in TBS containing 0.1% Tween-20 and incubated overnight at 4 °C with primary antibody diluted in blocking buffer. Blots were washed 3X with TBS containing 0.1% Tween-20 and incubated for 1 hr RT with secondary antibody conjugated to horseradish peroxidase diluted into blocking buffer (Jackson ImmunoResearch—West Grove, PA). Blots were washed a final 3X and imaged using Clarity Western ECL substrate (BioRad) and a BioRad Gel Doc XR+ system.

### Immunoprecipitation of mNeonGreen-PAVE1

Both endogenous alleles of PAVE1 were tagged at their N-termini with mNeonGreen (68) as described above. 3-8.0 x 10^8^ WT 427 control and mNeonGreen-PAVE1 cells were harvested and lysed in lysis buffer [10 mM Na3PO4 pH 7.4, 150 mM NaCl, 0.5% glycerol, 7% sucrose, and 0.5% NP-40 with 1 mM DTT; 1mM PMSF and 1X HALT (ThermoFisher) were added as protease inhibitors]. Cytoskeletal proteins were solubilized with sonication, after which the lysate was clarified by centrifugation at 17.1K *x g* for 20 min at 4 °C. Cleared supernatant material was diluted by half to decrease the concentration of detergent and incubated with magnetic beads conjugated to the camelid nanobody mNeonGreen-Trap (Chromotek—Munich, Germany) for 1 hr 4 °C. The beads were washed 6X with lysis buffer without glycerol, sucrose, and detergent. Beads were immediately processed for mass spectrometry, or bound proteins were eluted by adding SDS-PAGE lysis buffer and incubating the beads at 99 °C for 10 min. Eluted proteins were separated by SDS-PAGE and the gel was silver stained (ThermoFisher Scientific) for imaging.

### Mass spectrometry

#### On-beads sample preparation of control and mNeonGreen-PAVE1 immunoprecipitates

The supernatants from mNeonGreen-PAVE1 and corresponding 427 WT control immunoprecipitations were removed and replaced with 100 μL 0.1 M ammonium bicarbonate in water. Cys residues were reduced and alkylated using 5 mM dithiothreitol in 0.1 M ammonium bicarbonate for 1.5 hr, and 10 mM iodoacetamide in 0.1 M ammonium for 45 min in the dark, respectively. Both reactions proceeded at 37°C in a thermal mixer. A 16 hr, 37°C on-beads tryptic digestion was performed on a thermal mixer with constant agitation using a 1:20 (w/w) enzyme:protein ratio for Promega modified sequencing grade trypsin. Tryptic peptides were then removed and the digestion was quenched by the addition of concentrated formic acid to yield a final pH of 3. Proteolytic peptides were desalted using Pierce™ C18 Spin Columns (ThermoFisher Scientific) per manufacturer’s instruction, dried to completion, resuspended in 0.1% formic acid in water (Solvent A) and quantified using a Thermo Scientific Nanodrop Spectrophotometer.

#### Mass spectrometry-based proteomics analysis

mNeonGreen-PAVE1 and WT 427 control peptide sample concentrations were matched using Solvent A prior to injection onto a Waters 75 μm x 250 mm BEH C18 analytical column using a Thermo Scientific Ultimate 3000 RSLCnano ultra-high performance liquid chromatography system directly coupled to a Thermo Scientific Q Exactive HF Orbitrap mass spectrometer. A 1 hr, linear reversed-phase gradient (Solvent B: 0.1% formic acid in acetonitrile) at 300 nL/min flow was used to separate peptides prior to mass analysis.

Peptides were ionized directly into the Q Exactive HF via nanoflow electrospray ionization. High energy collision dissociation (HCD) MS/MS peptide interrogation was achieved using a top 15 data-dependent acquisition method in positive ESI mode. All raw data were searched against the Uniprot *T. brucei* reference proteome (identifier AUP000008524, database accessed 09/11/2018 and updated 04/27/2018) plus the full mNeonGreen-PAVE1 protein sequence using the MaxQuant software suite (69). Search parameters included the following: 1% false discovery rate cutoff at the protein and peptide-spectrum match levels, trypsin cleavage specificity with 2 missed cleavages, 5 amino acids/peptide minimum, and the “LFQ” algorithm was used to provide label-free quantitation. All other MaxQuant parameters were kept at default values. Search output files were compiled into Scaffold Q+S (Proteome Software, Inc.) for data visualization and further analysis.

### Recombinant Expression and Purification of the PAVE Complex

For recombinant expression in bacteria, the entire coding sequences of PAVE1 and PAVE2 were amplified from *T. brucei* genomic DNA. The solubility-enhancing maltose-binding protein (MBP) was fused to an oligo-His tag followed by a TEV protease site and appended to the N-terminus of mNeonGreen-PAVE1. The MBP-oligoHis-TEV-mNeonGreen-PAVE1 construct was inserted into a MalpET expression vector containing kanamycin resistance. The Strep-tag (70) peptide sequence was appended to the N-terminus of PAVE2 and inserted into a pOKD5 expression vector containing ampicillin resistance. Both vectors were co-transformed into BL21 (DE3) bacteria. 0.5 L of cells were grown to mid-log phase (OD_600_ ~0.6) and overexpression of the proteins were induced by adding 0.4 mM IPTG at 16 °C overnight. Cells were collected by centrifugation and lysed in lysis buffer [100 mM Tris pH 8.0, 300 mM NaCl, 0.5% NP-40, 0.5 mM DTT, with the addition of 1 mM PMSF and 1X HALT]. After sonication, the lysate was clarified by centrifugation at 17K *x g* and the supernatant was batch-incubated for 45 min at 4 °C with 3 mL of Ni-NTA agarose resin that had been pre-treated with 0.5 M imidazole pH 7.4 followed by equilibration with lysis buffer (ThermoFisher Scientific). The protein-bound resin was passed over a column and washed with 20 column volumes of Wash10 buffer [100 mM Tris pH 8.0, 300 mM NaCl, 0.5 mM DTT, 10 mM imidazole] followed by 20 column volumes of Wash30 buffer [100 mM Tris pH 8.0, 300 mM NaCl, 0.5 mM DTT, 30 mM imidazole]. Bound proteins were eluted in 1 mL fractions using elution buffer [100 mM Tris pH 8.0, 300 mM NaCl, 0.5 mM DTT, 200 mM imidazole]. Peak fractions were pooled and dialyzed overnight at 4 °C into Strep buffer [100 mM Tris pH 8.0, 150 mM NaCl, 1 mM EDTA]. Dialyzed fractions were treated with TEV protease (1:25 molar ratio) to remove the MBP-oligoHis-TEV fusion from mNeonGreen-PAVE1 for 5 hr RT, after which the complex was passed over Strep-Tactin XT resin (IBA Lifesciences—Goettingen, Germany). Bound complex was eluted using Buffer BXT containing 50 mM biotin (IBA Lifesciences). Protein samples were diluted into SDS-PAGE lysis buffer and incubated at 99 °C for 10 min for separation by SDS-PAGE and Coomassie staining. Purified PAVE complex was then dialyzed into storage buffer [100 mM Tris pH 8.0, 300 mM NaCl, 1 mM DTT, 50% glycerol] overnight at 4 °C. This preparation typically yielded ~200 μg of total protein that was stable for several months at −20 °C.

### In vitro Microtubule Interaction Assays

Microtubules were polymerized from cycled bovine tubulin (PurSolutions—Nashville, TN) mixed with Cy5-labled bovine tubulin (PurSolutions) and stabilized with 10 μM Taxol (Cytoskeleton Inc.—Denver, CO) as previously described (71), (42). The PAVE complex was clarified to remove aggregates by ultracentrifugation at 400K *x g* at 2 °C.

#### In-solution microtubule binding experiments

5-500 nM PAVE complex was mixed with microtubules diluted in blocking buffer [10 mM imidazole pH 7.4, 50 mM KCl, 1 mM EGTA, 4 mM MgCl2 supplemented with 2 mg mL^-1^ k-casein, 0.1% Plurionic F-68 (ThermoFisher Scientific), 10 μM Taxol, and an oxygen-scavenging system (3 mg ml^-1^ glucose, 0.1 mg ml^-1^ glucose oxidase and 0.18 mg ml^-1^ catalase)] and incubated for 20 min RT. Flow cells were constructed from glass coverslips and treated with rigor-like kinesin to attach microtubules to the glass surface as previously described (42,71). The flow cells were then blocked with two passages of blocking buffer. The complex-microtubule mixture was passed over flow cells, and any complex-microtubule mixture not captured by the rigor-like kinesin was washed away with two passages of blocking buffer. Flow cells were imaged using epifluorescence as above. Figures are representative of three independent biological replicates using two separate PAVE complex preparations.

#### Total internal reflection fluorescence microscopy experiments

PEGylated flow cells were prepared to minimize background as previously described (71). Rigor-like kinesin was passed over the PEGylated flow cell, followed by two passages of blocking buffer. For the PAVE complex binding experiment, microtubules diluted in blocking buffer were allowed to attach to the rigor-like kinesin on the PEGylated surface, after which unbound microtubules were washed away with blocking buffer. 5 nM PAVE complex was added to the flow cell and immediately imaged using TIRF microscopy. For the PAVE complex patch stability experiment, PAVE complex patches were pre-seeded on microtubules in solution for 10 min RT and passed through the flow cell. Unbound material was removed with three washes of blocking buffer, after which the flow cell was imaged using TIRF microscopy. Figures and movies are representative of three independent biological replicates using two separate PAVE complex preparations.

#### Subtilisin-treated microtubule binding assay

The C-terminal tails of Taxol-stabilized Cy5 microtubules were cleaved using Subtilisin A (Sigma Aldrich) as previously described (72). A 1:15 molar ratio of microtubules to subtilisin A were diluted into BRB80 blocking buffer [80 mM PIPES pH 6.8 with KOH, 1 mM MgCl_2_, 1 mM EGTA, supplemented with 2 mg mL^-1^ k-casein, 0.1% Plurionic F-68, 10 μM Taxol, and an oxygen-scavenging system as above] and incubated for 45 min at 30 °C. The cleaving reaction was stopped with the addition of 10 mM PMSF for 10 min at 30 °C. Samples of the microtubules were diluted into SDS-PAGE lysis buffer and incubated at 99 °C for 10 min for western blotting to confirm tail cleavage as above. Control or subtilisin-treated microtubules were incubated with 5 nM or 50 nM PAVE complex for 20 min RT and passed over prepared glass flow cells and imaged using epifluorescence as described above. Figures are representative of four independent biological replicates using one PAVE complex preparation.

---

**Supplementary Figure 1.**
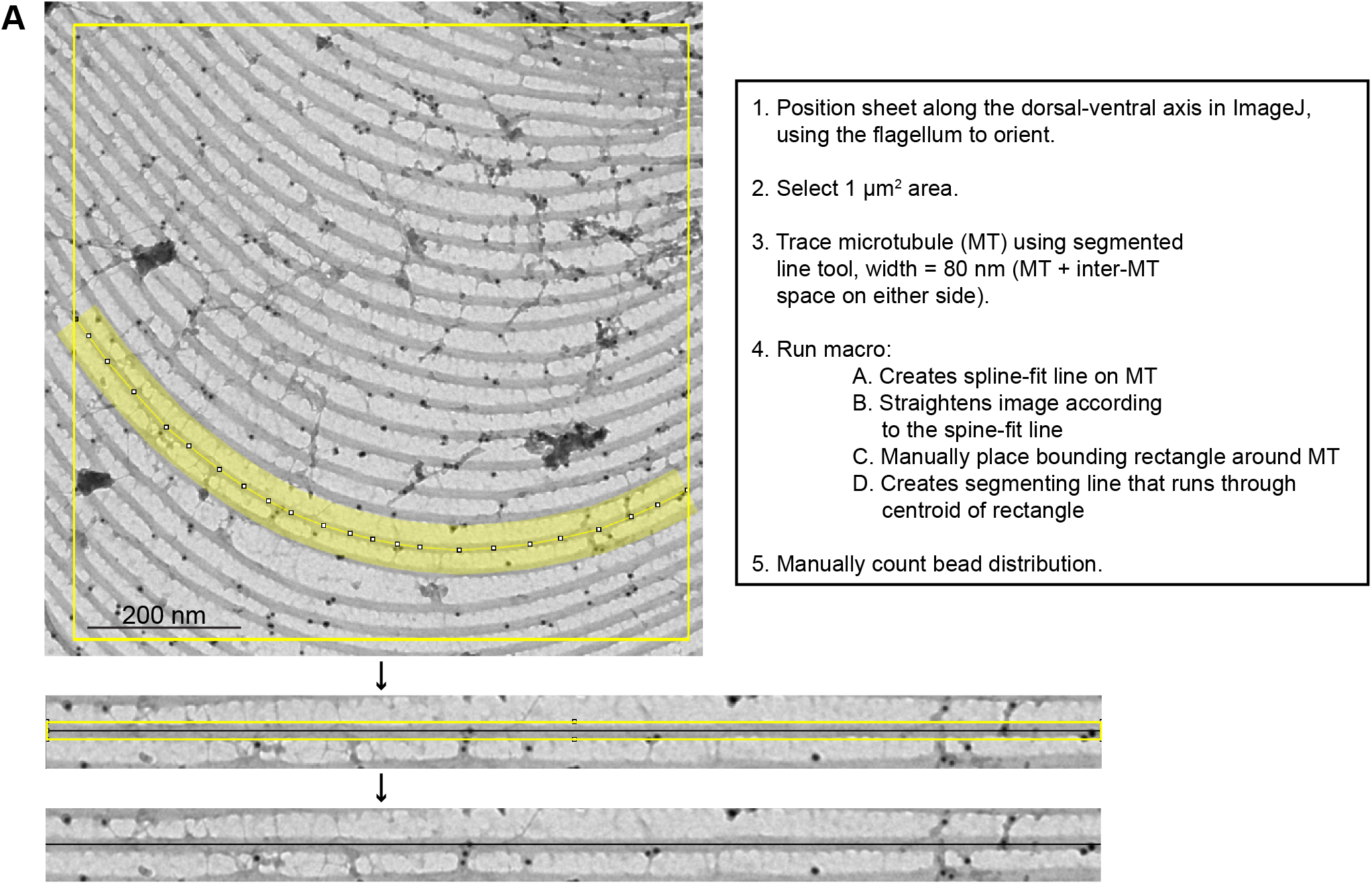
Quantification methodology used to classify immunogold label distribution in subpellicular array sheets. **(A)** A semi-automated macro was written in ImageJ to facilitate the classification gold bead distribution as dorsal, centroid, or ventral, as well as cross-link associated, along individual microtubules of subpellicular array sheets.

**Supplementary Figure 2.**
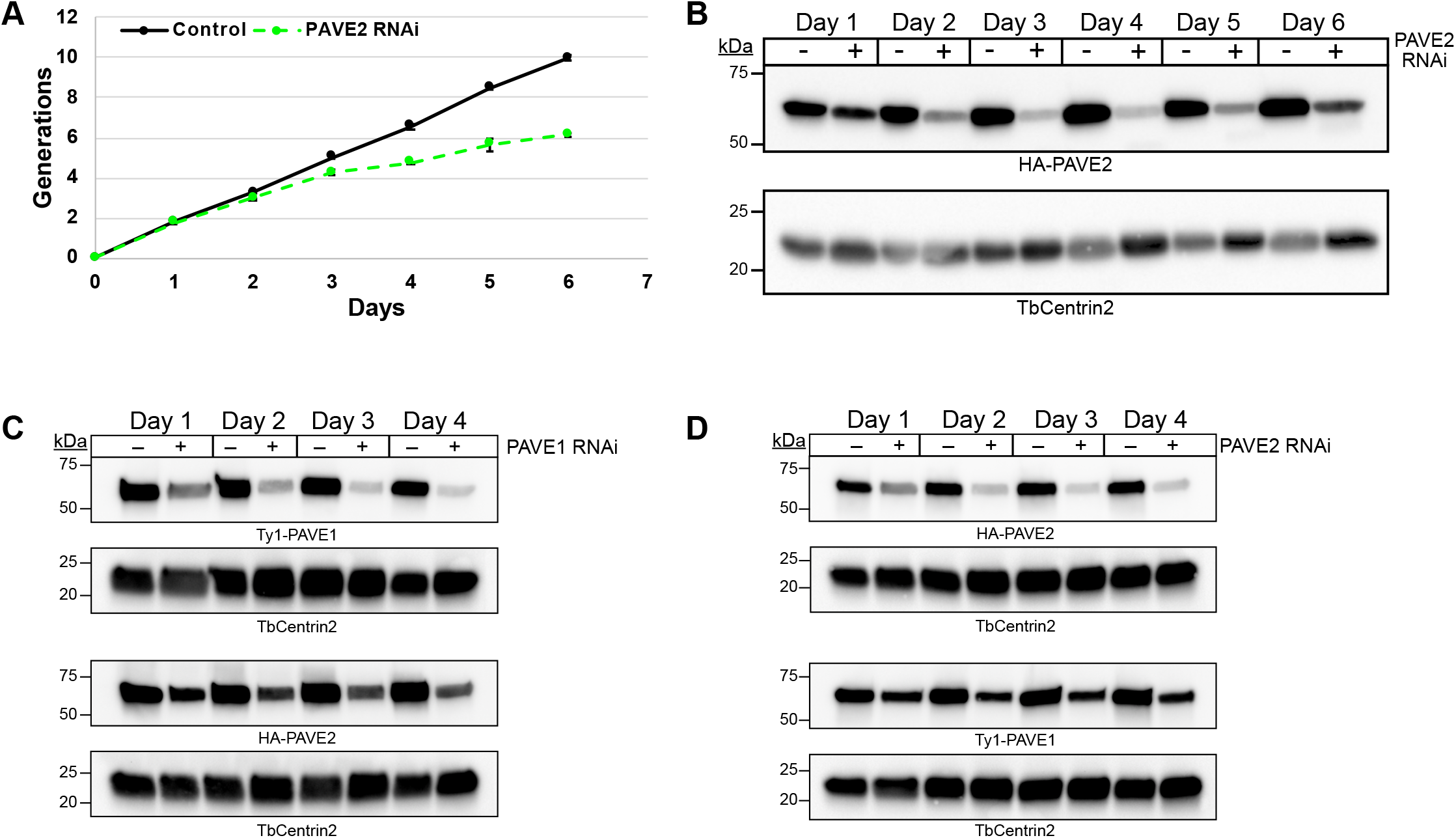
PAVE1 and PAVE2 require each other for stability in vivo. **(A)** Generation plot of cell growth after induction of PAVE2 RNAi. The cell density was monitored every 24 h in control and PAVE2 RNAi conditions. The curve is the mean of N=3 independent biological replicates. **(B)** Lysates were collected every 24 h from control and PAVE2 RNAi cells during 6 d of PAVE2 depletion. Lysates were western blotted with antibodies against HA and TbCentrin2 as a loading control. **(C)** Lysates from control and PAVE1 RNAi cells in which PAVE2 is endogenously tagged were collected for western blotting every 24 h during PAVE1 depletion. Duplicate lysates were western blotted, as PAVE1 and PAVE2 are the same molecular weight. **(D)** Lysates from control and PAVE2 RNAi cells in which PAVE1 is endogenously tagged were collected for western blotting every 24 h during PAVE2 depletion. The lysates were blotted in duplicates as in (C).

**Supplementary Figure 3.**
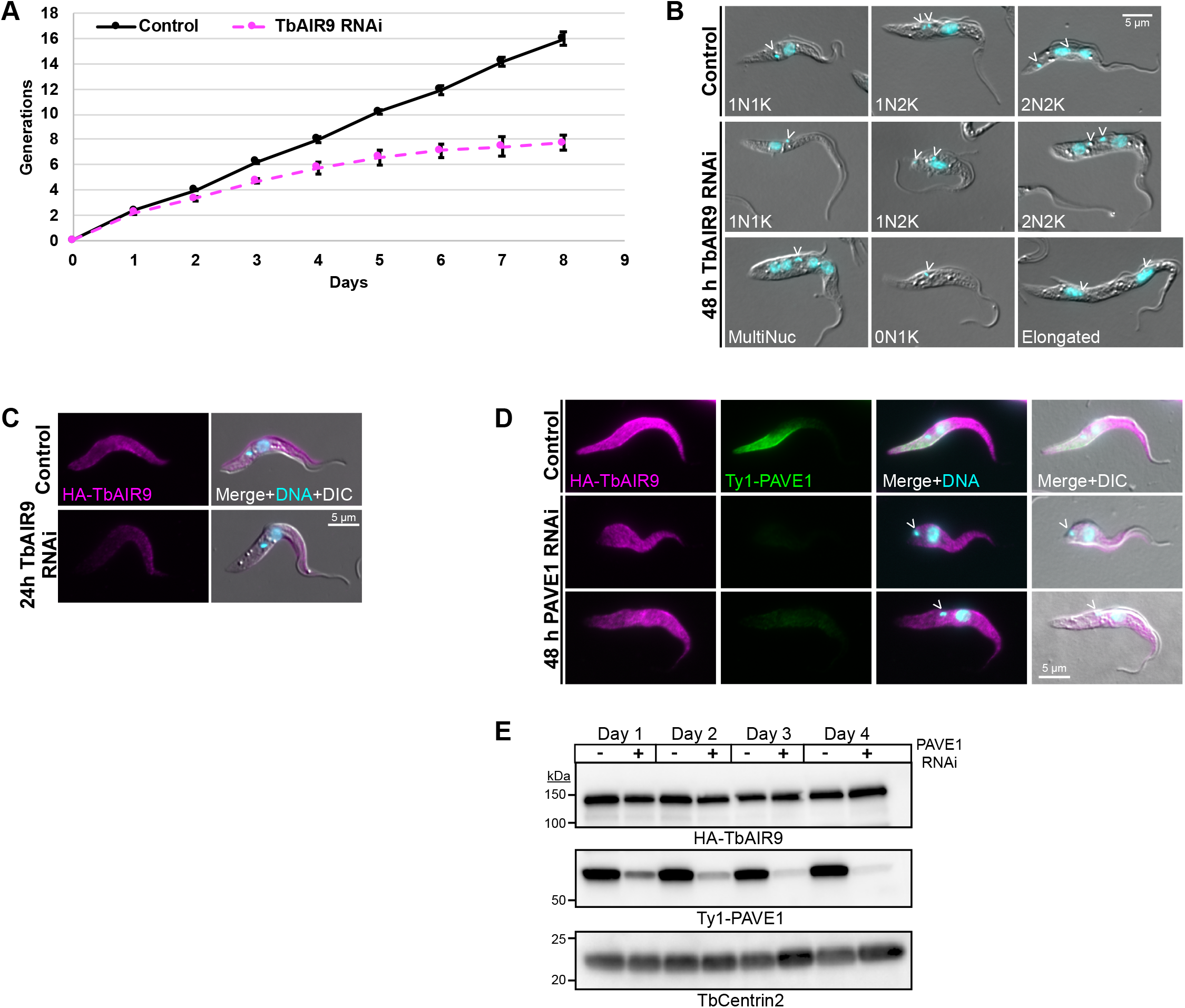
TbAIR9 does not require PAVE1 for localization or stability in vivo. **(A)** Generation plot of cell growth after induction of TbAIR9 RNAi. Cell concentration was monitored in control and TbAIR9 RNAi cells every 24 h. The curve is the mean of N=3 independent biological replicates. **(B)** Control and TbAIR9 RNAi cells were fixed in paraformaldehyde after 2 d of TbAIR9 depletion. Images are representative morphologies in control and TbAIR9 RNAi conditions. The epimastigote-like morphotype and elongated cell bodies were common in cells depleted of TbAIR9, as previously reported [(34)]. Open arrowheads indicate the kinetoplast. **(C)** Control and TbAIR9 RNAi cells were fixed with paraformaldehyde and stained after 24 h of TbAIR9 depletion. TbAIR9 localization initially disappears from the cell posterior during TbAIR9 RNAi, as previously reported. **(D)** PAVE1 RNAi was induced in cells in which TbAIR9 is endogenously tagged with the HA epitope tag. Cells were collected for fixation in paraformaldehyde after 2 d of PAVE1 depletion and stained for epifluoresence microscopy. The localization pattern of TbAIR9 does not change after PAVE1 depletion. **(E)** Lysates were collected from control and PAVE1 RNAi cells in which TbAIR9 is endogenously tagged every 24 h during PAVE1 depletion and western blotted. TbAIR9 stability is not affected by PAVE1 depletion.

**Supplementary Figure 4.**
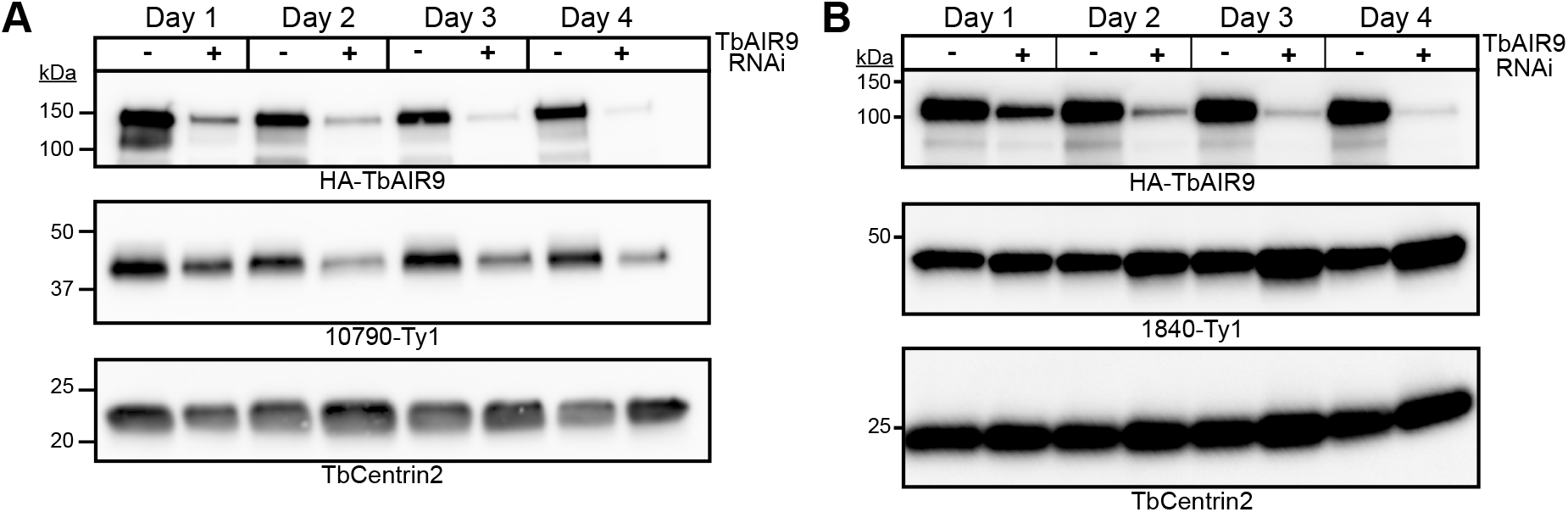
TbAIR9 depletion has differing effects on protein stability of array-associated proteins. **(A)** Lysates were collected from control and TbAIR9 RNAi cells in which Tb927.9.10790 was endogenously tagged every 24 h during TbAIR9 depletion and western blotted. 10790-Ty1 protein levels decrease during TbAIR9 depletion. **(B)** Lysates were collected from control and TbAIR9 RNAi cells in which Tb927.11.1840 was endogenously tagged every 24 h during TbAIR9 depletion and western blotted. 1840-Ty1 protein levels are stable and increase during TbAIR9 depletion.

**Movie 1.** TIRF microscopy of 5 nM PAVE complex binding to Taxol-stabilized Cy5-labeled bovine microtubules. Images were taken every 10 sec for 20 min. Movie is 10 frames per second.

**Movie 2.** TIRF microscopy of 5 nM PAVE complex patches pre-seeded for 10 min on Taxol-stabilized Cy5-labeled bovine microtubules. Unbound protein was washed out prior to imaging. Images were taken every 10 sec for 20 min. Movie is 10 frames per second.

## ACKNOWLEDGEMENTS

The authors would like to thank Dr. Jeremy Balsbaugh (University of Connecticut Proteomics and Metabolomics Facility) for the mass spectrometry analysis. We would also like to thank Geoff Williams (Brown University Leduc Bioimaging Facility) for helpful advice on our TEM preparations. We thank Dr. Sebastian Lourido (Massachusetts Institute of Technology) for the Ty1 antibody and Dr. Jack Sunter (Oxford Brookes University) for the TAT1 antibody. We thank Dr. Alexandra Deaconescu (Brown University) for the pOKD5 plasmid. We would like to thank TrypTag, whose data helped us identify Tb927.9.10790 and Tb927.11.1840. TrypTag is a data resource generated through a Wellcome Trust biomedical resource grant [108445/Z/15/Z] that was awarded to Prof. Keith Gull (University of Oxford), Prof. Mark Carrington (University of Cambridge), Prof. Christiane Hertz-Fowler (Wellcome Trust), Dr. Richard Wheeler (University of Oxford), Dr. Jack Sunter (Oxford Brookes University), Dr. Samuel Dean (University of Warwick) and Prof. Sue Vaughan (Oxford Brookes University). Work in C.L.d.G.’s laboratory is supported by the National Institutes of Health (F31AI143163 to A.N.S. and AI112953 to C.L.d.G.).

## Abbreviations

FAZ: Flagellum attachment zone
MAP: microtubule associated protein
N: nucleus
K: kinetoplast
iEM: immunogold electron microscopy
TEM: transmission electron microscopy
RNAi: RNA interference
mNG: mNeonGreen
MBP: maltose binding protein
TIRF: total internal reflection fluorescence microscopy

